# Uptake-independent killing of macrophages by extracellular aggregates of *Mycobacterium tuberculosis* is ESX-1 and PDIM-dependent

**DOI:** 10.1101/2023.01.11.523669

**Authors:** Chiara Toniolo, Neeraj Dhar, John D. McKinney

## Abstract

*Mycobacterium tuberculosis* (*Mtb*) infection is initiated by inhalation of small numbers of bacteria into lung alveoli, where they are phagocytosed by resident macrophages. Intracellular replication of *Mtb* leads to death of the infected macrophages, release of bacterial aggregates, and rapid growth of the extracellular aggregates on host-cell debris. Here, we show that extracellular *Mtb* aggregates can evade phagocytosis by killing macrophages in a contact-dependent but uptake-independent manner. We use single-cell time-lapse fluorescence microscopy to show that contact with extracellular *Mtb* aggregates triggers macrophage plasma membrane perturbation, cytoplasmic calcium accumulation, and pyroptotic cell death. These effects depend on the *Mtb* type VII secretion system ESX-1, however, this system alone cannot induce calcium accumulation and macrophage death in the absence of the *Mtb* surface-exposed lipid phthiocerol dimycocerosate. Unexpectedly, we found that ESX-1-mediated secretion of the EsxA/EsxB virulence factors is not required for uptake-independent killing of macrophages after contact with extracellular *Mtb* aggregates. In the absence of EsxA/EsxB secretion, killing is mediated by the 50-kDa isoform of the ESX-1-secreted protein EspB, while blocking secretion of both EsxA/EsxB and processed EspB reduces killing to background levels. Treatment with a small-molecule ESX-1 inhibitor reduces uptake-independent killing of macrophages by *Mtb* aggregates, suggesting that novel therapies targeting this anti-phagocytic mechanism could prevent the propagation of extracellular bacteria within the lung.

**Significance statement:** *Mycobacterium tuberculosis* (*Mtb*) can survive inside the lung macrophages that normally provide the first line of defense against bacterial infections. Intracellular replication of *Mtb* ultimately results in the death and lysis of infected macrophages, allowing the bacteria to spread to other cells and propagate the infection. Our study shows that extracellular *Mtb* aggregates that form on the debris of dead host cells can induce macrophage death in a contact-dependent but uptake-independent manner, allowing the bacteria to evade the host defenses associated with uptake by macrophages. Killing of macrophages by extracellular *Mtb* aggregates is driven by the *Mtb* ESX-1 secretion system and the surface-exposed lipid phthiocerol dimycocerosate. Our results suggest that novel drugs targeting *Mtb* factors required for host-cell killing by extracellular *Mtb* aggregates may reduce bacterial spreading and expansion of necrotic tuberculosis lesions, which are known to be poorly penetrated by conventional antibiotics.

## Introduction

The success of *Mycobacterium tuberculosis* (*Mtb*) as a human pathogen hinges on the ability to survive attacks by the immune cells that mediate host defenses against lung infections. Tuberculosis infections are initiated by the inhalation and implantation of small numbers of bacteria in the lung alveoli, where they are rapidly phagocytosed by resident alveolar macrophages (1, 2). Upon phagocytosis, *Mtb* uses different strategies to adapt to the intracellular milieu, such as prevention of phagolysosome acidification (3), metabolic adaptation (4), lysis of the phagosomal membrane, and escape into the cytoplasm (5, 6). Replication of cytoplasmic bacteria ultimately leads to lysis of the infected macrophage, which is a key process in spreading the infection to other cells (7).

Lysis of infected macrophages *in vitro* results in rapid replication of *Mtb* on the debris of the dead cells and formation of extracellular bacterial aggregates (8). Formation of *Mtb* bacterial aggregates is also observed *in vivo* in animal models of infection (9–11) and in human patients (12–18), and necrotic foci in the lungs are typically associated with high burden of extracellular bacteria (9, 12, 14). Phagocytosis of extracellular *Mtb* aggregates by newly-recruited macrophages triggers cycles of intracellular infection and host-cell lysis, contributing to progressive propagation of the bacteria (19, 20). At early stages of tuberculosis, immune cells are recruited to the site of infection by the pro-inflammatory signaling associated with host-cell death (7, 21). At later stages, host-cell death in granulomas, the multi-cellular structures that encapsulate the bacteria and limit their propagation, can lead to caseation, cavitation, and dissemination of the infection to the airways, allowing transmission of *Mtb* to other hosts (22). Despite the importance of these processes in tuberculosis pathogenesis, the host-cell pathways that mediate cell death are still controversial and different forms of cell death have been reported (23), including apoptosis (24, 25), necrosis (26, 27), necroptosis (28, 29), pyroptosis (30), and other non-canonical forms of death mediated by type I interferon (31) or tumor necrosis factor signaling (32).

Induction of host-cell death by intracellular bacteria requires the *Mtb* ESX-1 type VII secretion system (5, 6). This system secretes several proteins, including the EsxA/EsxB heterodimer, EspA, EspB and EspC, which are required for *Mtb* virulence (33). In particular, the ESX-1-secreted EsxA protein has been shown to mediate the breakdown of phagolysosomal membrane integrity and translocation of *Mtb* into the cytoplasm (5, 6). Phagolysosomal membrane rupture also requires the complex lipid phthiocerol dimycocerosate (PDIM) displayed on the bacterial cell surface (25, 33). Rupture of the phagolysosome or cytoplasmic sensing of *Mtb* DNA can activate several different death pathways in the host cells (28, 32, 34), while physical contact between cytoplasmic bacteria and the inner face of the host-cell plasma membrane can lead to plasma membrane damage and death by pyroptosis (30). EsxA has also been shown to be required for induction of macrophage death after phagocytosis of extracellular *Mtb* growing on the debris of dead host cells (19).

Killing of host cells by intracellular bacteria may promote tuberculosis pathogenesis by allowing *Mtb* to evade macrophage defense mechanisms (35) and to grow rapidly on the debris of dead host cells (8). As an alternative to killing of macrophages by intracellular bacteria, we considered the possibility that evasion of host-cell defenses might also be achieved by killing of macrophages by extracellular bacteria in an uptake-independent manner. Previous studies have shown that intracellular *Mtb* can induce plasma membrane damage and inhibit plasma membrane repair in infected macrophages (30, 36). Contact with extracellular *Mtb* also induces host-cell plasma membrane damage and inflammasome activation, but it was not reported whether these events resulted in host-cell death (30). In an earlier study, contact of macrophages with extracellular *Mtb* did not lead to cell death when uptake was inhibited with cytochalasin D treatment (26). In both studies, macrophages were exposed to single-cell bacterial suspensions rather than aggregates of *Mtb* formed after death of infected macrophages and extracellular growth on host-cell debris.

In this study, we use time-lapse fluorescence microscopy to investigate the dynamic interaction between macrophages and *Mtb* at the single-cell level. We show that contact of macrophages with extracellular aggregates of *Mtb* results in pyroptotic host-cell death in an uptake-independent manner. Killing of macrophages by extracellular *Mtb* is dependent on aggregation *per se*, because similar numbers of non-aggregated individual bacteria are less efficient at inducing macrophage death upon contact. Before dying, macrophages exhibit signs of local plasma membrane perturbation at the site of contact with *Mtb* aggregates and progressive accumulation of calcium in the cytoplasm. These two processes require the *Mtb* ESX-1 secretion system and the surface-exposed lipid phthiocerol dimycocerosate (PDIM). Unexpectedly, we found that secretion and proteolytic processing of EspB is responsible for driving plasma membrane perturbation, calcium accumulation, and host-cell death in the absence of EsxA/EsxB secretion, revealing previously unknown roles for this protein in *Mtb* pathogenesis.

## Results

### *M. tuberculosis* aggregates induce contact-dependent, uptake-independent killing of macrophages

We used time-lapse fluorescence microscopy to track individual mouse bone marrow-derived macrophages (BMDMs) infected with *Mycobacterium tuberculosis* (*Mtb*) expressing tdTomato (movie S1). After phagocytosis, intracellular *Mtb* replicates (figure S1A) and eventually kills and lyses host cells (figure S1B; movies S1-2). Once released from lysed macrophages, *Mtb* replicates rapidly on the debris of the dead cells to form extracellular aggregates (figure S1A,C; movies S1-2). As previously described (19, 20), we observed that macrophages that interact with these bacterial aggregates eventually die (figures 1A; S2A; S3A), leading to uncontrolled bacterial proliferation on the host-cell debris and formation of large extracellular *Mtb* aggregates (movie S1). Bacterial aggregates from axenic cultures induce macrophage death with comparable timing (figure 1B,C; movie S3), demonstrating that neither a previous intracellular passage of the bacteria in macrophages nor the presence of cellular debris on the aggregates is required to induce macrophage death. Physical interaction between bacterial aggregates and macrophages is required to induce death of the infected cells, as highlighted by the observation that bystander macrophages near infected cells show the same survival as uninfected cells over the course of the infection (figures 1D; S3A).

**Figure 1.**
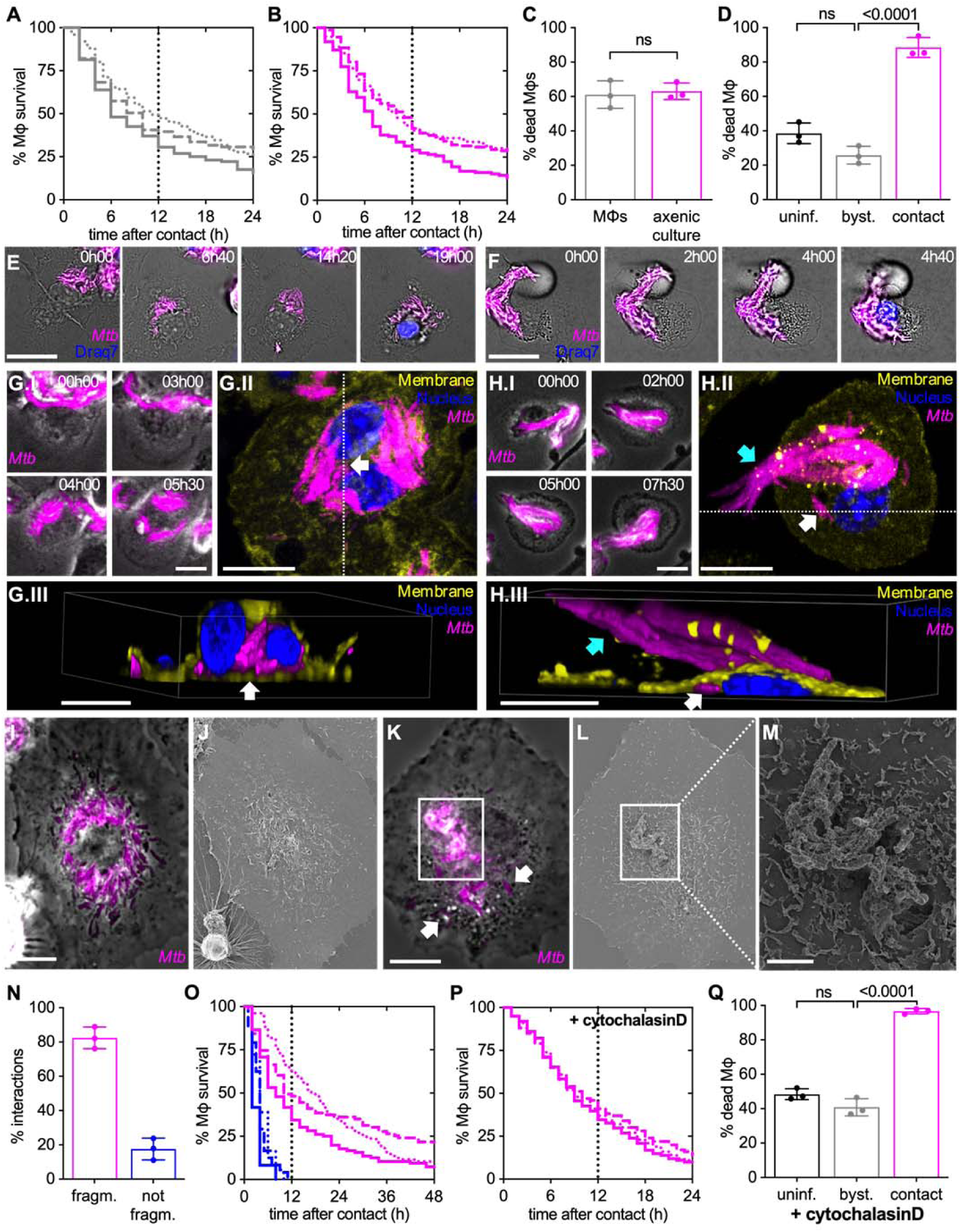
Aggregates of *Mtb* kill macrophages in a contact-dependent but uptake-independent manner. **(A,C,N,O)** Mouse bone marrow-derived macrophages (BMDMs) infected with *Mtb* Erd-tdTomato and imaged by time-lapse microscopy at 1- or 2-hour intervals for 132 hours. **(B-M)** BMDMs infected with aggregates of *Mtb* Erd-tdTomato from axenic culture and imaged by time-lapse microscopy at 1-hour intervals for 60 hours. **(A-B)** Percentage of macrophages surviving after interaction with an extracellular *Mtb* aggregate originating from the debris of dead macrophages (A) or from axanic culture (B). Each line represents an independent experimental replicate (n = 3 replicates with ≥ 100 cells per replicate). **(C)** Percentage of macrophages that die within the first 12 hours after stable contact with an *Mtb* aggregate originating either from a dead macrophage (MΦ) or from an axenic culture. **(D)** Percentage of macrophages that die by the end of the experiment (by 60 h post infection). Macrophages in “contact” represents the fraction of cells that interact with an *Mtb* aggregate during the course of the experiment. We define as “bystander” (byst.) the macrophages that are in the same sample of the infected ones but do not establish physical contact with an *Mtb* aggregate during the course of the experiment. “Uninfected” (uninf.) macrophages are not exposed to *Mtb*. **(E)** Example of a macrophage that interacts with an *Mtb* aggregate (magenta) (0h00), fragments it (3h00), redistributes the bacteria in a “bullseye” pattern around the nucleus (12h00), and dies (16h00). **(F)** Example of a macrophage that stably interacts with an *Mtb* aggregate without fragmenting it (0h00-09h00) and ultimately dies (12h00). In **E** and **F**, Draq7 staining of the nucleus (in blue) is used as a marker for cell death. Scale bars, 20 μm. **(G-H)** BMDMs infected with aggregates of *Mtb* were imaged by time-lapse microscopy (every 30 min for up to 13.5h) followed by fixation, immunostaining and imaging by confocal microscopy. *Mtb* in magenta, nuclei stained with Hoechst (blue), membrane staining with anti-CD45 antibody in yellow. White arrows indicate intracellular bacteria, cyan arrows indicate extracellular bacteria. All scale bars, 10 μM. **(G.I,H.I)** Time-lapse microscopy image-series of macrophages that interacts with *Mtb* aggregates and fragment (G) or do not fragment (H) them. **(G.II,H.II)** Max intensity projection of confocal microscopy images of the same macrophages shown in panels I. **(G.III,H.III)** 3-D reconstruction of the cells imaged in panels II, images are cropped in x or y in the position indicated by the white dotted lines in panels **II** to show the inside of the cell. **(I-M)** BMDMs infected with aggregates of *Mtb* were imaged by time-lapse fluorescence microscopy followed by SEM. **(I)** Example of a macrophage showing the typical “bullseye” pattern of bacterial redistribution around the cell nucleus. **(J)** Correlative SEM image of (I). **(K)** Example of a macrophage interacting with an extracellular *Mtb* aggregate without fragmentation and redistribution of bacteria. Time-lapse microscopy image-series of these macrophages are shown in figure S5. **(L-M)** Correlative SEM image of (K). Scale bars, 10 μm (I-L) or 2 μm (M). **(N)** Percentage of macrophage-*Mtb* aggregate interactions resulting (“fragm.”, pattern exemplified by panel E) or not resulting in fragmentation (“not fragm.”, pattern exemplified by panel F) of the *Mtb* aggregate. The quantification was performed by manual annotation of all the macrophage-*Mtb* aggregate interactions observed in the time-lapse microscopy movies. **(O)** Percentage of macrophages surviving over time after uptake and fragmentation of *Mtb* aggregates or after contact without fragmentation. Each line represents an independent experimental replicate. **(P-Q)** BMDMs were treated with cytochalasin D to prevent bacterial uptake, infected with aggregates of *Mtb* and imaged by time-lapse microscopy at 1-hour intervals for 60 hours. **(P)** Percentage survival over time for macrophages in contact with *Mtb* aggregates. Time 0 represents the time when stable contact with an aggregate begins. Each line represents an independent experimental replicate (n = 3 replicates with ≥ 100 cells per replicate). **(Q)** Percentage of cytochalasin D-treated macrophages that die by the end of the experiment (by 60 h post infection). “Bystander” and “in contact” and “uninfected” macrophages defined as in panel (D). In **(C,D,N,Q)** each symbol represents the percentage of events for a single experimental replicate (n ≥ 100 events per replicate). Bars represent means and standard deviations; *P* value calculated using a t test (C) or a one-way ANOVA test (D,Q), ns, *P* values > 0.05.

Time-lapse microscopy reveals two different patterns of long-term interaction of macrophages with *Mtb* aggregates. Approximately 80% of interactions between macrophages and extracellular *Mtb* aggregate result in fragmentation of the *Mtb* aggregate within 2-3 hours after first contact and redistribution of the bacteria in a “bullseye” pattern around the host-cell nucleus (figure 1E,N; movie S4). In the remaining ∼ 20% of interactions, macrophages remain in stable contact with extracellular *Mtb* aggregates for hours without any sign of fragmentation of the aggregate or perinuclear redistribution of bacteria (figure 1F,N; movie S5). We hypothesized that the first pattern follows uptake of the *Mtb* aggregate by the macrophage (figure 1E), whereas the second pattern represents a stable association of the *Mtb* aggregate with the macrophage without uptake (figure 1F). We tested this hypothesis by using correlative single-cell time-lapse microscopy followed by macrophage plasma membrane immunostaining and confocal imaging or by scanning electron microscopy (SEM). Bacterial aggregates that upon interaction with a macrophage get fragmented and acquire the “bullseye” pattern (figures 1G.I-II; S4A.I-II,C.I-II) are localized inside the macrophage (figures 1G.III; S4A.III,C.III) and are not visible on the surface of the macrophage in SEM images (figures 1I-J, S5A-C). Conversely, aggregates that do not undergo fragmentation and perinuclear redistribution (figures 1H.I-II; S4B.I-II,D.I-II) remain extracellular (figures 1H.III; S4B.III,D.III) and readily visible on the surface of the macrophage by SEM (figures 1K-M; S5A,D-I). Occasionally, some individual bacteria can be detached from the main extracellular aggregate visible on the surface of the cell and be internalized (figures 1HII-III,K; S4B.II-III,D.II-III, white arrows). These results are consistent with the notion that the first pattern represents intracellular bacterial aggregates whereas the second pattern represents extracellular bacterial aggregates.

Both these patterns are observed in macrophages that die upon interaction with a bacterial aggregate (figure 1E,F,N). Notably, macrophages that do not take up and fragment the *Mtb* aggregates invariably die within 12 hours of first contact with the aggregate (figure 1O), while macrophages that take up and fragment an *Mtb* aggregate survive longer (figure 1O). We thus asked whether extracellular *Mtb* aggregates may induce macrophage death prior to uptake. By treating macrophages with cytochalasin D, an actin polymerization inhibitor that blocks phagocytosis without affecting *Mtb* growth (figure S6), we confirmed that uptake is not required for killing of macrophages by extracellular *Mtb* aggregates (figures 1P; S2B; S3B; movie S6-7). We also observed that uninfected bystander macrophages treated with cytochalasin D survive over the same time course (figures 1Q; S3B; movie S6), further confirming that killing of macrophages by extracellular *Mtb* aggregates needs physical contact despite not requiring uptake.

### Bacterial aggregation is important for contact-dependent, uptake-independent killing of macrophages

Phagocytosis of single *Mtb* bacilli or aggregated *Mtb* induces macrophage death in a dose-dependent manner (figure S7A,B) (20). Interestingly, as previously reported (20), we observe that aggregated bacteria kill more efficiently than similar amounts of single bacilli (figure S7A,B) leading to faster bacterial propagation (figure S7C,D). We thus asked whether bacterial aggregation *per se* is also important for contact-dependent, uptake-independent killing of macrophages by *Mtb*. We found that macrophages treated with cytochalasin D to prevent bacterial uptake and challenged with increasing numbers of non-aggregated bacteria (figure 2A-E) are killed in a dose-dependent manner (figure 2F). However, comparisons of macrophages challenged with similar numbers of aggregated or non-aggregated bacteria (figure 2A-E; cf. “high” and “aggregate”) show that aggregated bacteria are significantly more toxic than non-aggregated bacteria (figure 2F). We conclude that both uptake-dependent and contact-dependent (uptake-independent) killing of macrophages by *Mtb* depends on both the number of bacteria and their aggregation status.

**Figure 2.**
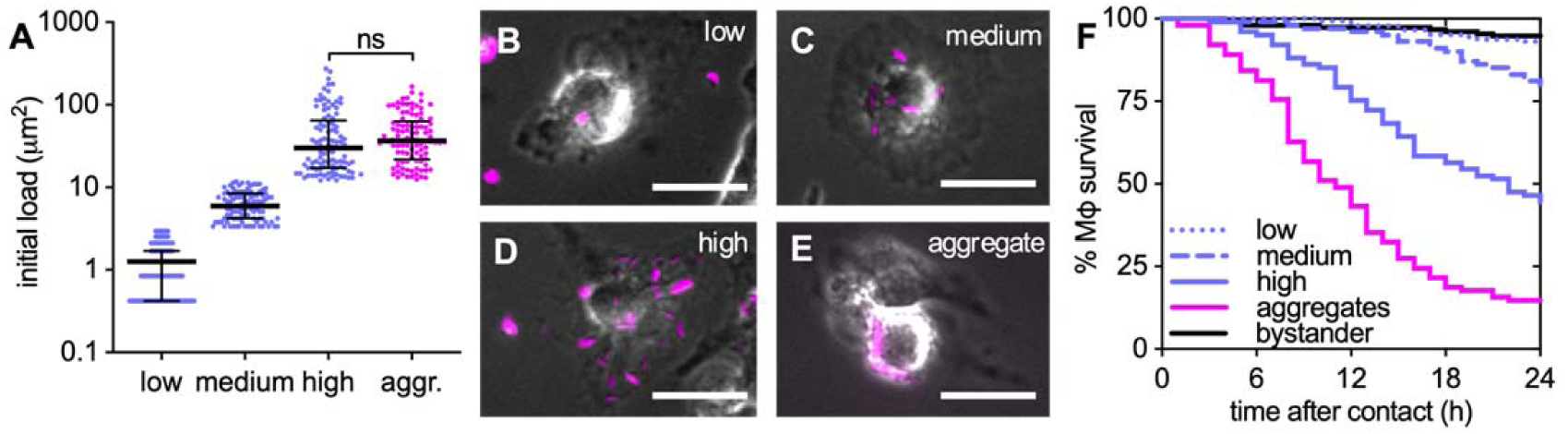
Bacterial aggregation is important for contact-dependent uptake-independent killing of macrophages. **(A-F)** BMDMs were treated with cytochalasin D and infected with aggregated or non-aggregated *Mtb* Erd-tdTomato and imaged by time-lapse microscopy at 1-hour intervals for 60 hours. **(A)** Initial bacterial load calculated as the total fluorescent area per macrophage. Infected individual macrophages are binned into low, medium and high initial load according to the number of single bacteria they are in contact with. For comparison, we selected macrophages in contact with aggregates (aggr.) of *Mtb* having a load similar to the high gate. Gates were set at <3.0 μm^2^ (low), 3.0-12.0 μm^2^ (medium), or >12.0 μm^2^ (high and aggregates) per macrophage (n ≥ 100 cells per gate). The area of one bacterium is included between 0.5 and 2 μm^2^. Each symbol represents the bacterial load of one macrophage. Bars represent median and interquartile range. P-value calculated using an unpaired Mann-Whitney test. **(B-E)** Examples of macrophages infected with low (B), medium (C), or high (D) doses of non-aggregated bacteria or with bacterial aggregates (E). Scale bars, 20 μm. **(F)** Percentage survival over time for macrophages in contact with increasing doses of non-aggregated bacteria or with bacterial aggregates. Time 0 for cells “in contact” represents the time when stable contact with the bacteria begins. Each line represents a biological replicate (n ≥ 100 cells per condition).

### Plasma membrane perturbation in macrophages at the site of contact with extracellular *Mtb* aggregates

We used single-cell time-lapse fluorescence microscopy to identify events leading up to the death of macrophages in contact with *Mtb* aggregates. Cytochalasin D-treated BMDMs establish stable interactions with extracellular *Mtb* aggregates despite being unable to internalize them (figures 3A; S8A,B). Staining with fluorescent Annexin V shows that approximately 75% of these macrophages (figure 3C), display Annexin V-positive plasma membrane domains at the site of contact with an *Mtb* aggregate, indicating the presence of exposed phosphatidylserine associated with a local plasma membrane perturbation (figures 3A; S8A-C; movies S7,S8). After appearance of the local Annexin V-positive membrane domain, affected macrophages survive for 11 ± 10 hours (figure 3D) and become Annexin V-positive over the entire plasma membrane only after death (figure 3A,B; S8D; movies S7,S8). Notably, the cells that develop local Annexin V-positive membrane domains seem to die faster than the cells that don’t show this pattern (figure 3D). Correlative SEM revealed that the membrane of cytochalasin D-treated Annexin V-negative macrophages extends and partially covers the associated bacterial aggregate (figure 3E,F). At later stages of the interaction, when the plasma membrane in contact with the *Mtb* aggregate becomes Annexin V-positive, the membrane undergoes fragmentation and blebbing (figure 3G). Aggregates that don’t contact macrophages never become Annexin V-positive (figure S8C). However, we observe Annexin V-positive foci colocalize not only with markers for the macrophage plasma membrane (figure S8A,B) but also with more distal areas of the bacterial aggregates that do not stain positive for plasma membrane markers (figure S8B). Moreover, we see that upon macrophage death, aggregates in contact with dead cells retain some Annexin V-positive material on their surface (figure S8C, movie S8). Vesicle budding and shedding is a common ESCRT III-mediated membrane repair strategy that allows removal of damaged portions of the plasma membrane and wound resealing (37). We therefore think that Annexin V-positive foci might represent both areas of damaged membrane as well as macrophage plasma membrane vesicles that were released and which stick to the hydrophobic surface of the bacterial aggregates. Interestingly, the fluorescence intensity of the Annexin V-positive domains did not increase over time, but we detected several single intensity peaks (movie S8), suggesting that multiple discrete damaging events happen upon interaction with a *Mtb* aggreate.

**Figure 3.**
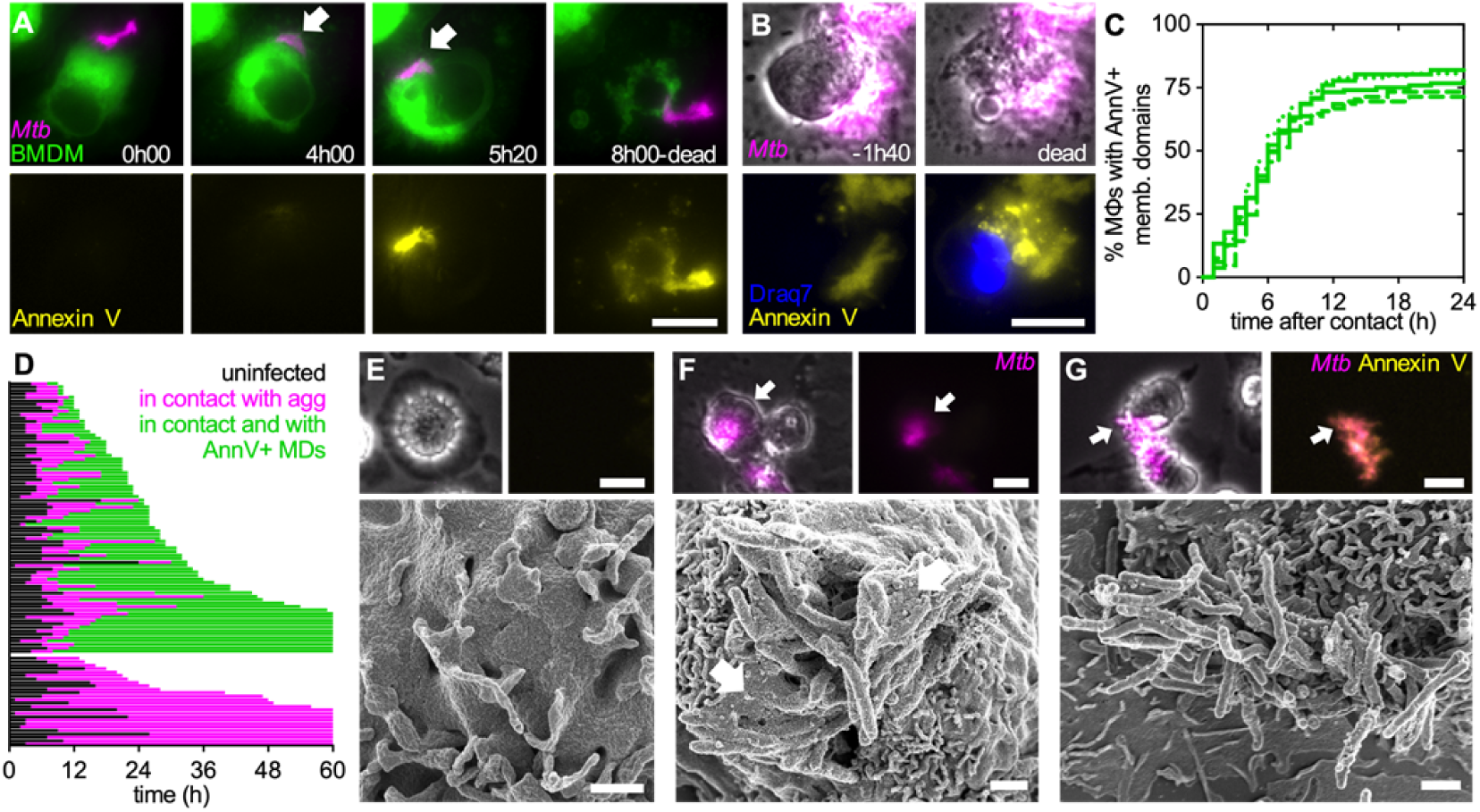
Uptake-independent killing of macrophages by *Mtb* aggregates involves intimate contact between the macrophage membrane and the bacterial aggregate. **(A,B)** BMDMs expressing a membrane-targeted tdTomato were treated with cytochalasin D, infected with aggregates of *Mtb* expressing GFP, and imaged by time-lapse microscopy at 20-minute intervals for 24 hours. **(A)** Example of a BMDM interacting with an extracellular *Mtb* aggregate. Top panels show the macrophage plasma membrane (green) and the bacterial aggregate (magenta). Intimate interaction between the macrophage plasma membrane and the bacterial aggregate begins at 4h00 (indicated by the white arrow). Bottom panels show the Annexin V-647 positive (yellow) plasma membrane domain that appears at 5 hours 20 minutes and co-localizes with the bacterial aggregate (magenta). At 8h00, plasma membrane integrity is lost (top panel) and Annexin V accumulates throughout the dead cell (bottom panel). Scale bar, 20 μm. **(B)** Example of a BMDM that dies after interacting with an extracellular *Mtb* aggregate. Top panels show a macrophage in contact with an extracellular *Mtb* aggregate (magenta) 1h40 before death (left) and just after death (right). Bottom panels show an Annexin V-FITC positive (yellow) plasma membrane domain; Draq7 (blue) stains the macrophage nucleus only after the cell dies. Scale bar, 10 μm. **(C-D)** Cytochalasin D-treated BMDMs were infected with aggregates of *Mtb* Erd*-*tdTomato in the presence of Annexin V-FITC and imaged by time-lapse microscopy at 1-hour intervals for 60 hours. **(C)** Percentage of macrophages in contact with *Mtb* aggregates that show local Annexin V-positive membrane domains at the site of contact with the bacteria over time. Time 0 represents the time when stable contact with an aggregate begins. Each line represents an independent experimental replicate (n = 5 replicates with ≥ 100 cells per replicate). **(D)** Behavior of 105 individual macrophages in contact with *Mtb* aggregates. Each line represents the life span of an individual cell; the fraction of the line in black represents the time spent as uninfected, the fraction of the line in magenta represents the time spent interacting with an *Mtb* aggregate and the fraction in green represents the time upon formation of local Annexin V-positive membrane domains (MDs) at the site of contact with the *Mtb* aggregate. **(E-G)** Cytochalasin D-treated BMDMs were infected with aggregates of *Mtb* Erd-tdTomato in the presence of Annexin V-FITC. Selected cells were imaged by time-lapse fluorescence microscopy followed by SEM. Arrows in top panels indicate the cell imaged by SEM. Top left: brightfield image of macrophage. Top left and right: fluorescence image of the bacterial aggregate (magenta). Top right: fluorescence image of Annexin V staining (yellow). Bottom: correlative SEM image. Scale bars, 20 μm in top panels and 1 μm in bottom panels. **(E)** Example of a bystander BMDM that is not in contact with a bacterial aggregate. **(F)** Example of an Annexin V-negative BMDM interacting with a *Mtb* aggregate. Arrows in bottom panel indicate areas of intact plasma membrane interacting with bacterial aggregates. **(G)** Example of a BMDM with blebbed Annexin V-positive membrane domains interacting with an *Mtb* aggregate.

These observations suggest that induction of macrophage death does not require uptake of the bacteria *per se* but does involve intimate contact between the macrophage membrane and the bacterial aggregate followed by local membrane perturbation.

### Contact with extracellular *Mtb* aggregates induces calcium accumulation in macrophages

After escaping from the phagosome into the cytoplasm of infected macrophages, intracellular *Mtb* has been shown to damage the plasma membrane and to affect its permeability to ions (30). We therefore asked whether the membrane perturbations observed at the site of contact between macrophages and extracellular *Mtb* aggregates are linked to aberrant ion permeability. We used a Ca^2+^-dependent fluorescent dye to monitor intracellular Ca^2+^ dynamics in cytochalasin D-treated BMDMs exposed to extracellular *Mtb* aggregates. Before death, cells interacting with *Mtb* aggregates accumulate significantly more cytoplasmic Ca^2+^ than uninfected bystander cells (figure 4A,B). Ca^2+^ also accumulates in cells interacting with *Mtb* aggregates that do not die over the course of the experiment (figure S9A,B).

**Figure 4.**
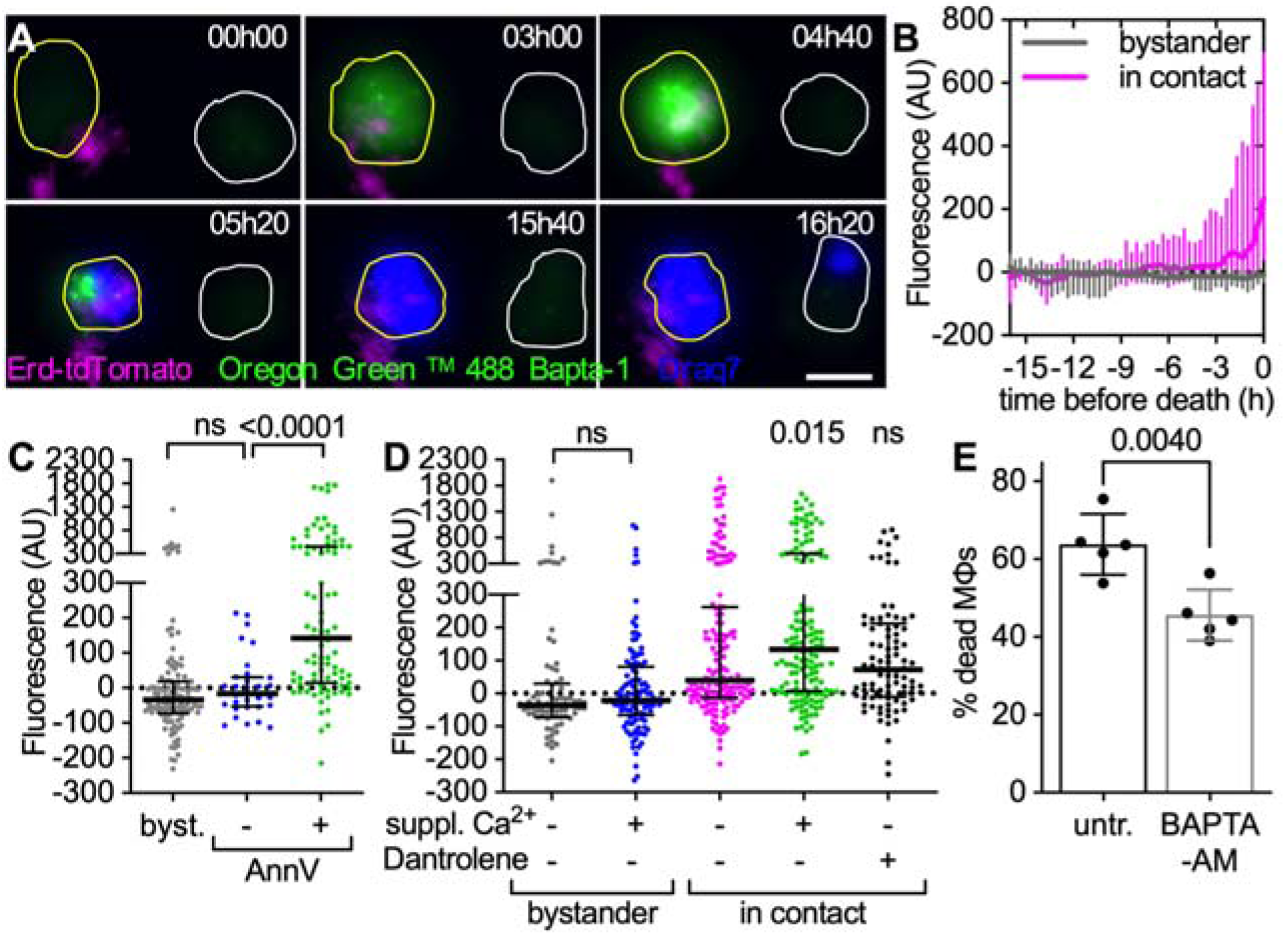
Extracellular *Mtb* aggregates induce cytoplasmic calcium accumulation in cytochalasin D-treated macrophages. **(A-D)** Cytochalasin D-treated BMDMs were infected with aggregates of *Mtb* Erdman WT, stained with the membrane-permeable dye Oregon Green 488 Bapta-1 AM to visualize intracellular Ca^2+^ and imaged by time-lapse microscopy at 20-minute intervals for 24 hours. Oregon Green 488 Bapta-1 AM fluorescence values at each time point were normalized to time 0 for uninfected bystander cells and to the time of first contact with an *Mtb* aggregate for infected cells. In (**C-D**) values for infected cells correspond to the time of death after first contact or 16 hours post-contact for cells that survive. Values for uninfected bystander cells correspond to the time of death or 16 hours. Each symbol represents a single macrophage. Black bars represent median and interquartile range. **(A)** Examples of BMDMs dying with (yellow outline) or without (white outline) interacting with extracellular *Mtb* aggregates (magenta). Cell outlines are based on brightfield images (not shown). Oregon Green 488 Bapta-1 AM fluorescence is shown in green. Cell death is indicated by Draq7 nuclear staining (blue). Scale bar, 20 μm. **(B)** Oregon Green 488 Bapta-1 AM fluorescence over time in dying bystander macrophages (n=15) and in dying macrophages in contact with an *Mtb* aggregates (n=63). Lines represent median fluorescence values for all cells, error bars represent interquartile ranges.The distributions of the fluorescence values at time of death in bystander and dying macrophages are significantly different, *p* value <0.0001, calculated using a Welch’s t test. **(C)** Oregon Green 488 Bapta-1 AM fluorescence for uninfected bystander cells (byst.; n=121) and for infected cells with (+; n=91) or without (–; n=32) Annexin V-positive plasma membrane domains at the site of contact with an *Mtb* aggregate. *P*-values were calculated using a Krustal-Wallis test; ns, *P* values > 0.05. **(D)** Oregon Green 488 Bapta-1 AM fluorescence in bystander and infected macrophages incubated in medium with or without 10 mM Ca^2+^ and dantrolene. *P*-value for bystander cells calculated using an unpaired Mann-Whitney test. *P*-values for infected cells calculated using a Krustal-Wallis test comparing the treated samples to the untreated control; ns, *P* values > 0.05. (n=81, 120, 165, 152, 94, respectively). **(E)** BMDMs treated with cytochalasin D were infected with aggregates of *Mtb* Erd-tdTomato and imaged by time-lapse microscopy at 1-hour intervals for 60 hours. Percentage of macrophages that die within the first 12 hours after stable contact with an *Mtb* aggregate without (untr.) or with supplementation of BAPTA-AM. Each symbol represents the percentage of dead macrophages for a single experimental replicate (n ≥ 60 cells per replicate). Bars represent means and standard deviations, *P* value calculated using a t test.

Cytoplasmic Ca^2+^ accumulation is significantly higher in cells with Annexin V-positive membrane domains at the site of contact with *Mtb* aggregates, suggesting that local membrane perturbations could make the cells permeable to small ions such as Ca^2+^ (figures 4C; S9C,D). Appearance of Annexin V-positive local membrane domains and intracellular Ca^2+^ accumulation was also observed in BMDMs upon contact with extracellular *Mtb* aggregates in the absence of cytochalasin D (figure S10A-D), thereby validating the use of cytochalasin D treatment to study uptake-independent killing of macrophages by extracellular *Mtb* aggregates.

Cells incubated in medium supplemented with Ca^2+^ accumulate more intracellular Ca^2+^ upon contact with *Mtb* aggregates in comparison to cells incubated with regular medium (figure 4D). Moreover, macrophages treated with dantrolene, a RyR inhibitor that inhibits Ca^2+^ release from the endoplasmic reticulum, still accumulate Ca^2+^ in the cytoplasm (figure 4D). Ca^2+^ chelation with the cell-permeant chelator BAPTA-AM significantly reduces the percentage of macrophages that die upon contact with an extracellular *Mtb* aggregate (figure 4E) without affecting formation of Annexin V-positive membrane domains (figure S11). These observations suggest that *Mtb*-induced plasma membrane perturbation may cause influx of Ca^2+^ from the extracellular milieu into the cytoplasm, leading to cell death.

### Extracellular *Mtb* aggregates induce uptake-independent pyroptosis in macrophages

Perturbations in intracellular calcium homeostasis have been linked to different cell death pathways (32, 38–40). We asked whether uptake-independent killing of macrophages by extracellular *Mtb* aggregates is mediated by one of the known cell death pathways. We observed that uninfected bystander macrophages that die show signs of Caspase-8 activation (figures 5A; S12C,D), a common marker induced in apoptosis (figure S12A,B) (38), but rarely display “specks” of apoptosis-associated speck-like protein containing a CARD (ASC) (figure 5C), a marker for inflammasome activation induced in pyroptosis (figure S12E,F) (41). Conversely, macrophages that die upon contact with extracellular *Mtb* aggregates usually display ASC-specks (figure 5B,C) but seldom stain positive for cleaved Caspase-8 (figure 5A). These observations are consistent with a previous study showing that contact between extracellular *Mtb* and plasma membrane can induce inflammasome activation (30) and suggest that contact with extracellular aggregates of *Mtb* induces a death pathway different from apoptosis. Cytochalasin D-treated macrophages in contact with *Mtb* aggregates also show Ca^2+^-dependent enhanced activation of Caspase-1 (figures 5D; S12G-K), an effector caspase that is autoproteolytically activated by the inflammasome complex and that promotes pyroptotic cell death by inducing cleavage of gasdermin D (GSDMD) (40). Macrophages in contact with *Mtb* aggregates do not show any enhanced activation of the necroptosis effectors RIP3 (figures 5E; S12L-P) or MLKL (figures 5F; S12Q-U). Taken together, these observations suggest that contact with extracellular *Mtb* aggregates may induce pyroptotic cell death in macrophages. In agreement with this interpretation, we observed that two different pyroptosis inhibitors (the NLRP3 inhibitor MCC950 and the Caspase-1 inhibitor VX765) block Caspase-1 activation (figure 5G) and partially reduce cell death (figure 5H) in cytochalasin D-treated macrophages in contact with *Mtb* aggregates, whereas inhibition of Caspase-8 or RIP3 has no effect (figure 5H). Notably, treatment with pyroptosis inhibitors does not affect *Mtb* growth (figure S13A), formation of Annexin V-positive membrane domains (figure S13B) or Ca^2+^ accumulation (figure S13C) in macrophages in contact with *Mtb* aggregates, suggesting that the Ca^2+^ influx depends on contact with the bacteria and not on GSDMD pore formation downstream of Caspase-1 activation.

**Figure 5.**
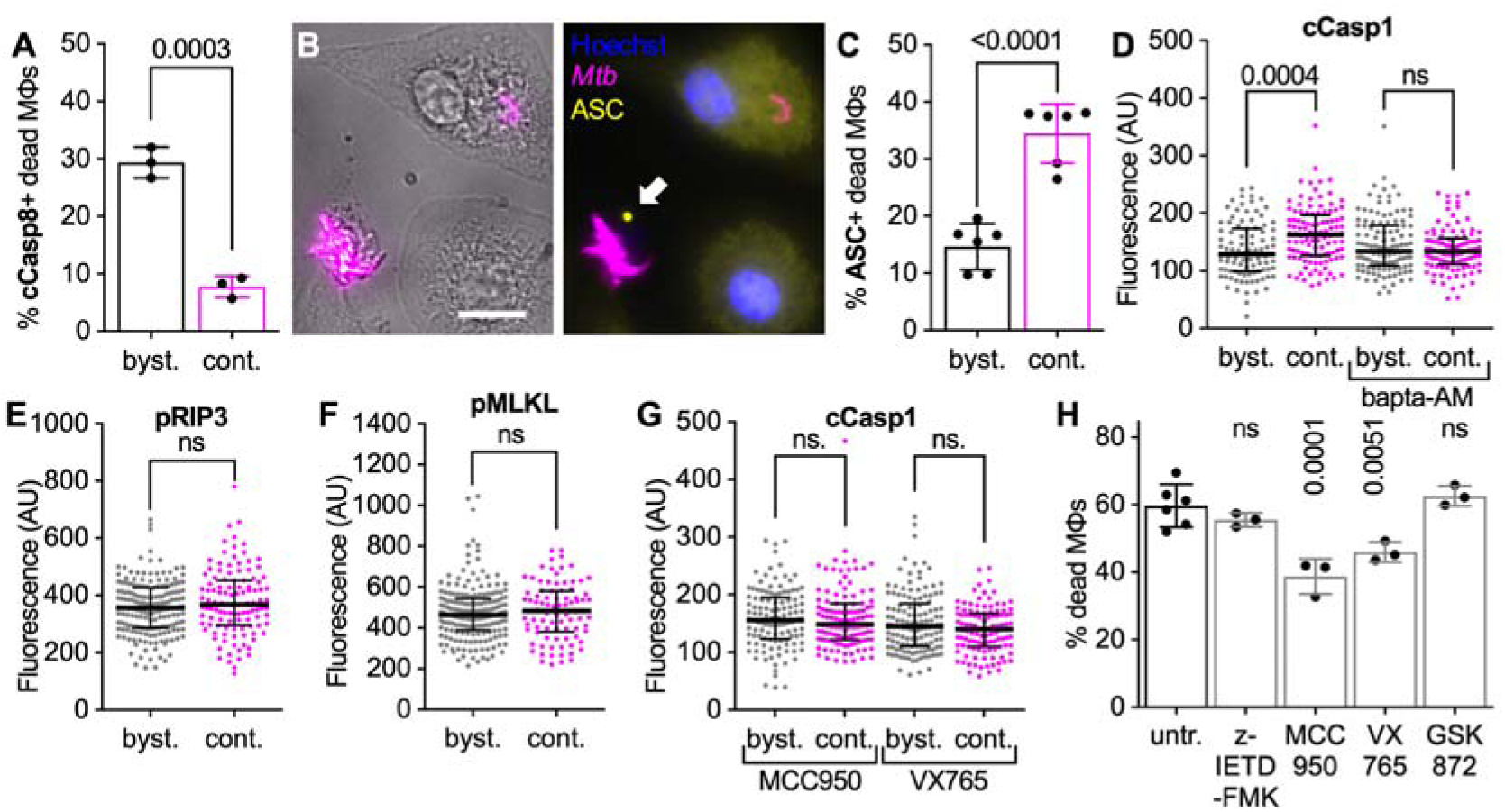
Extracellular *Mtb* aggregates induce inflammasome activation and pyroptosis in cytochalasin D-treated macrophages. **(A-G)** BMDMs treated with cytochalasin D were infected with aggregates of *Mtb* Erd-tdTomato, fixed at 24 hours post infection and processed for immunofluorescence with antibodies targeting cellular markers for cell death pathways. Controls and representative microscopy images for all the antibodies used provided in figure S5. Macrophages are defined as “in contact” when the body of a macrophage identified in brightfield images overlaps with a bacterial aggregate identified in the fluorescence channel (see figure S5 for representative examples). **(A)** Percentage of dead uninfected bystander macrophages (byst.) or dead macrophages in contact (cont.) with an extracellular *Mtb* aggregate that stain positive for cleaved Caspase-8 (cCasp8). **(B)** Representative example of a dead macrophage in contact with an extracellular *Mtb* aggregate (magenta) that display a ASC-specks (yellow, white arrow). Nuclei stained with Hoechst (blue). **(C)** Percentage of dead uninfected bystander macrophages (byst.) or dead macrophages in contact with an extracellular *Mtb* aggregate (cont.) that display ASC-specks. **(A,C)** Each symbol represents the percentage of positive dead macrophages for a single experimental replicate (n ≥ 80 cells per replicate). Bars represent means and standard deviations. *P* values were calculated using a t test. **(D-G)** Median fluorescence values for bystander macrophages (byst.) or macrophages in contact with an extracellular *Mtb* aggregate (cont.) stained with an **(D,G)** anti-cleaved Caspase-1 antibody (cCasp1), an **(E)** anti-phosphorylated RIP3 antibody (pRIP3) or **(F)** anti-phosphorylated MLKL antibody (pMLKL). In **(D)** BMDMs were untreated or incubated with Bapta-AM during the course of the infection. In **(G)** BMDMs were treated with MCC950 (NLRP3 inhibitor) or with VX765 (Caspase-1 inhibitor) during the course of the infection. Each symbol represents a single macrophage. Black bars represent median and interquartile range. P-value were calculated using an unpaired Mann-Whitney test; ns, *P* values > 0.05. (**D:** n=93, 108, 115, 101 respectively; **E:** n=182, 112 respectively; **F:** n=186, 84 respectively; **G:** n=121, 117, 129, 111 respectively). **(H)** BMDMs treated with cytochalasin D were infected with aggregates of *Mtb* and imaged by time-lapse microscopy at 1-hour intervals for 60 hours. Percentage of macrophages that die within the first 12 hours after stable contact with an *Mtb* aggregate. Macrophages were treated with the apoptosis inhibitor Z-IETD-FMK (Caspase-8 inhibitor); with pyroptosis inhibitors MCC950 (NLRP3 inhibitor) or VX765 (Caspase-1 inhibitor); or with necroptosis inhibitor GSK872 (RIP3 inhibitor). Each symbol represents the percentage of dead macrophages for a single experimental replicate (n ≥ 100 cells per replicate). Bars represent means and standard deviations. *P* values were calculated using a one-way ANOVA test comparing treated samples with the untreated control; ns, *P* values > 0.05.

### ESX-1 and PDIM are both required for uptake-independent killing of macrophages by *Mtb* aggregates

The *Mtb* ESX-1 type VII secretion system and the surface-exposed lipid phthiocerol dimycocerosate (PDIM) are required for phagosomal membrane damage, bacterial translocation into the cytoplasm, and induction of host-cell death by intracellular bacteria (42, 5, 43, 25). We therefore assessed whether these virulence factors also have a role in uptake-independent killing of macrophages by extracellular *Mtb* aggregates.

We found that a strain of *Mtb* with a large deletion in the ESX-1 operon (ΔRD1) (figure S14A) (42) does not induce uptake-independent killing of macrophages (figure 6A). Killing is largely restored by genetic complementation of the ΔRD1 mutant (figure 6A), confirming the essential role of the ESX-1 system. We also observed that a PDIM-deficient strain of *Mtb* with a disrupted *fadD26* gene (figure S15) does not induce uptake-independent macrophage death (figure 6B). PDIM production and ESX-1-dependent secretion have been suggested to be interdependent (44), however, in line with previous observations (25), we observed that the *fadD26* mutant show unaltered secretion of the ESX-1 secreted proteins (figures S14B,G; S17) and that the ΔRD1 strain is not impaired in PDIM production (figure S15), suggesting that mutations in these genes do not affect each other. Interestingly, when phagocytosis is not inhibited by cytochalasin D treatment, lack of PDIM has only a minor effect on macrophage killing by *Mtb* aggregates (figure S16), suggesting that this factor may have a less significant role in inducing uptake-dependent macrophage death.

**Figure 6.**
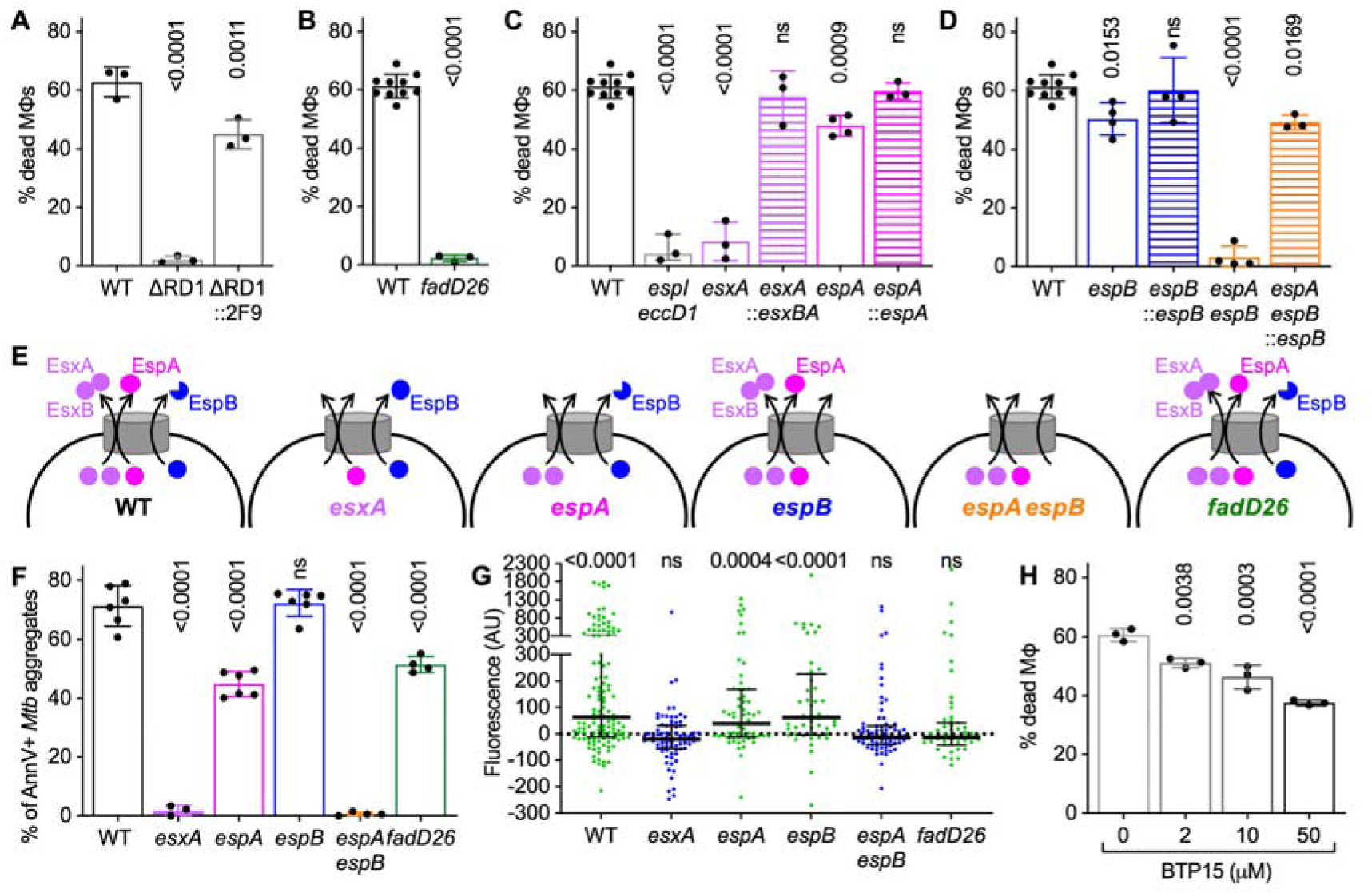
ESX-1-secreted proteins and PDIM are required for uptake-independent killing of macrophages by *Mtb* aggregates. **(A-D, H)** BMDMs treated with cytochalasin D were infected with aggregates of different *Mtb* strains and imaged by time-lapse microscopy at 1-hour intervals for 60 hours. The plots represent the percentage of macrophages that die within the first 12 hours after stable contact with an *Mtb* aggregate. Each symbol represents the percentage of dead macrophages for a single experimental replicate (n ≥ 3 replicates with ≥ 70 cells per replicate). Bars represent means and standard deviations. *P*-values were calculated using a one-way ANOVA test (**A**,**C**,**D,H**) or t test (**B**) comparing each strain to the wild-type reference strain (**A-D**) or the treated vs. untreated samples (**H**); ns, *P* values > 0.05. **(A)** Macrophages in contact with aggregates of *Mtb* H37Rv wild-type (WT), ΔRD1 mutant and complemented ΔRD1 mutant (ΔRD1::2F9). **(B)** Macrophages in contact with aggregates of *Mtb* Erdman wild-type (WT) or PDIM-deficient (*fadD26*) strains. **(C)** Macrophages in contact with aggregates of *Mtb* Erdman wild-type (WT) or mutant strains (*espI eccD1*, *esxA*, *espA*) and the complemented strains (*esxA*::*esxBA*, *espA*::*espA*). **(D)** Macrophages in contact with aggregates of *Mtb* Erdman wild-type (WT) or mutant strains (*espB*, *espA espB*) and the *espB*-complemented strains (*espB*::*espB*, *espA espB*::*espB*). **(E)** Schematic representation of EsxA, EsxB, EspA, EspB secretion pattern in the mutant strains used in this study (based on Western Blot and quantitative proteomics data shown in figures S5, S8 and S9). **(F)** BMDMs treated with cytochalasin D were infected with aggregates of different *Mtb* strains, incubated with Annexin V and imaged by time-lapse microscopy at 1-hour intervals for 48 hours. Percentage of macrophages that show Annexin V - positive membrane domains within the first 12 hours after entering in contact with *Mtb* aggregates. Each symbol represents a single experimental replicate (n ≥ 3 replicates with ≥ 90 cells per replicate). Bars represent means and standard deviations. *P*-values were calculated using a one-way ANOVA test. **(G)** Cytochalasin D-treated BMDMs were infected with aggregates of *Mtb* Erdman WT, stained with the membrane-permeable dye Oregon Green 488 Bapta-1 AM to visualize cytoplasmic Ca^2+^ and imaged by time-lapse microscopy at 20-minute intervals for 24 hours. Oregon Green 488 Bapta-1 AM fluorescence values at each time point were normalized to the time of first contact with an *Mtb* aggregate for infected cells. Values in the plot correspond to the time of death after first contact or 16 hours post-contact for cells that survive. Each symbol represents a single macrophage. Black bars represent median and interquartile range. P-value were calculated using a one sample Wilcoxon test; ns, *P* values > 0.05. (n=123, 64, 48, 71, 78, 55 respectively). **(H)** Percentage of macrophages that die within the first 12 hours after interaction with an *Mtb* aggregate. Infected cells are incubated with different concentration of BTP15 (0, 2, 10, 50 μM) during the course of the experiment.

### Expression but not secretion of EsxA/EsxB is required for uptake-independent killing of macrophages by *Mtb* aggregates

We used a panel of *Mtb* mutant strains to investigate how different ESX-1 components are involved in induction of uptake-independent macrophage death. We observed that macrophages in contact with aggregates of an *espI-eccD_1_* mutant with disrupted structural components of the ESX-1 secretion system (45) (figure S14B,C,E) do not die (figure 6C), demonstrating that a functional ESX-1 system is required for uptake-independent killing of macrophages.

Among the ESX-1 secreted proteins, EsxA has been shown to mediate breakdown of the phagolysosomal membrane (5). EsxA is co-secreted in a 1:1 heterodimer with EsxB (46, 47) but uncoupled secretion of these proteins can occur in the presence of an aberrant ESX-1 system (48, 49). We confirmed that in our hands, loss of EsxA eliminates EsxB expression and secretion (47, 50, 25) (figures S14B,C,F; S18A). We found that deletion of *esxA* reduces killing to nearly background levels (figure 6C) and that complementation with both *esxA* and *esxB* restores secretion of EsxB (figure S14F) and uptake-independent killing of macrophages (figure 6C). EsxA secretion requires EspA, another ESX-1-secreted protein (51), and deletion of *espA* abolishes EsxA/EsxB *secretion* (figures S14G; S18B) without affecting their *expression* (figure S14B,G) (52). Unexpectedly, we found that deletion of *espA* has only a slight impact on macrophage killing (figure 6C), suggesting that although *expression* of EsxA/EsxB is required for uptake-independent killing of macrophages, EspA-mediated *secretion* of these proteins is not required.

### EspB mediates uptake-independent killing in the absence of EsxA/EsxB secretion

The unexpected finding that uptake-independent killing of macrophages is eliminated by deletion of *esxA* but not *espA* is not linked to any discernable differences in the mutant strains’ production of PDIM (figure S15), aggregate morphology (figure S19A,B,F,G), or growth rates (figure S19H). However, quantitative proteomics revealed that deletion of *esxA* also decreases the secretion of EspK and EspB, two ESX-1-secreted proteins whose secretion is not affected by deletion of *espA* (figure S18A,B). In particular, deletion of *esxA* eliminates secretion of the cleaved 50-kDa isoform of EspB while having little or no impact on production and secretion of the full-length 60-kDa isoform (figure S14F). We therefore asked whether *Mtb* requires EspK or EspB to induce uptake-independent killing of macrophages.

We excluded the involvement of EspK by showing that an *espK* mutant strain induces uptake-independent killing of macrophages at levels comparable to wild-type *Mtb* (figure S20). On the other hand, we observed that although deletion of *espB* has only a slight impact on uptake-independent killing (figure 6D), similar to deletion of *espA* (figure 6C), deletion of both *espA* and *espB* reduces killing to background levels (figure 6D), similar to the impact of *esxA* deletion (figure 6C). The *espB* and *espA espB* mutants show no alteration in PDIM production (figure S15), aggregate morphology (figure S19C,D) or growth rates (figure S19H) and complementation with *espB* (figures S14H) restores macrophage killing to similar levels as the wild-type and *espA* strains (figure 6D). Quantitative proteomics revealed that *espB* deletion does not detectably affect the secretion of any other protein in either the wild-type or *espA* background (figure S18C,D), indicating that the impact of *espB* on killing is not due to loss of another secreted protein. We conclude that secreted EsxA/EsxB and EspB are both involved in uptake-independent killing of macrophages by *Mtb* aggregates because blocking secretion of only one of these proteins reduces but does not eliminate cell death.

### ESX-1 but not PDIM is required for membrane perturbation in macrophages contacting *Mtb* aggregates

Since macrophage killing requires both ESX-1 and PDIM, we asked whether these factors are also involved in triggering local membrane perturbations in macrophages after contact with *Mtb* aggregates. We found that aggregates of bacteria that secrete either EsxA/EsxB (*espB* mutant) or the 50-kDa isoform of EspB (*espA* mutant) are able to induce local membrane perturbations in contacted macrophages at nearly wild-type levels (figure 6F). In sharp contrast, blocking secretion of both EsxA/EsxB and the 50-kDa isoform of EspB (*esxA* and *espA espB* mutants) reduces membrane perturbations in contacted macrophages to nearly background levels (figure 6F). Although aggregates of PDIM-deficient *fadD26* bacteria do not kill macrophages (figure 6B), they still induce local membrane perturbations (figure 6F), consistent with the observation that this strain expresses and secretes EsxA/EsxB and EspB normally (figures S14B,G; S17). These results indicate that membrane perturbation *per se* is not sufficient to cause cell death and further support our conclusion that secreted EsxA/EsxB and EspB have overlapping roles in uptake-independent killing of macrophages.

### ESX-1 and PDIM are both required for calcium accumulation in macrophages contacting *Mtb* aggregates

Since membrane perturbation and calcium accumulation may be causally related, we evaluated the role of ESX-1 and PDIM in triggering calcium accumulation in macrophages after contact with extracellular *Mtb* aggregates. We found that aggregates of *Mtb* strains that do not induce membrane perturbations (*esxA* and *espA espB* mutants) also do not induce calcium accumulation in macrophages (figure 6G). However, we also found that PDIM-deficient bacteria (*fadD26* mutant) also fail to induce calcium accumulation in contacted macrophages (figure 6G) despite inducing membrane perturbation at nearly normal levels (figure 6F). Taken together, our results suggest that ESX-1-mediated membrane perturbation is not sufficient to induce calcium accumulation and macrophage death in the absence of PDIM.

### A small-molecule inhibitor of ESX-1 reduces uptake-independent killing of macrophages by *Mtb* aggregates

We asked whether BTP15, a small molecule that inhibits secretion of components of the ESX-1 system (53), can inhibit the uptake-independent killing of macrophages by *Mtb* aggregates. Treatment with BTP15 reduces secretion of EsxB and EspB (in particular the 50-kDa isoform) and uptake-independent killing of macrophages in a dose dependent manner (figures S14A-B; 6H). *Mtb* aggregates treated with BTP15 for 48 hours before infection, induce reduced uptake-independent killing of macrophages, even upon drug washout, excluding a possible direct effect of BTP15 on macrophages (figure S14I). We show that by inhibiting secretion of components of the ESX-1 system, BTP15 significantly reduces macrophage membrane perturbation (figure S21H), uptake-independent (figure 6H) and uptake-dependent (figure S21I) macrophage killing, without affecting bacterial growth or aggregation (figure S21D-G). These observations suggest that inhibitors targeting factors involved in macrophage killing by extracellular *Mtb* aggregates could interfere with bacterial evasion of host defense mechanisms that depend on phagocytic uptake.

## Discussion

Death of infected host macrophages is an important process in *Mtb* infections, as it promotes spreading of the intracellular bacteria to other host cells, inflammation, recruitment of immune cells, caseation, and disruption of granulomas (7, 21, 22). Induction of host-cell death by intracellular *Mtb* has been extensively investigated (23). Here, we extend these observations by using live-cell time-lapse microscopy of *Mtb*-infected macrophages and chemical inhibition of phagocytosis to demonstrate that *Mtb* can also induce macrophage death “from the outside” of the cell in a contact-dependent but uptake-independent manner. This experimental approach allowed us to focus only on death events induced by extracellular bacteria and to characterize this process, which may allow *Mtb* to evade stresses associated with phagocytic uptake by macrophages.

*Mtb* was previously shown to induce contact-dependent hemolysis of erythrocytes (54). However, the unusual structure and membrane composition of these anucleate non-phagocytic cells makes it difficult to extrapolate these results to phagocytic immune cells such as macrophages. Other cytotoxicity assays commonly used to quantify viability of host cells interacting with *Mtb* do not make a clear distinction between killing by extracellular vs. intracellular bacteria (55, 42, 52, 53). In macrophages treated with cytochalasin D to inhibit phagocytosis, contact with extracellular *Mtb* was shown to be sufficient to induce inflammasome activation but cell death was not reported (30). Here, we show that aggregates of extracellular *Mtb* are much more efficient in killing cytochalasin D-treated macrophages compared to similar numbers of single (non-aggregated) bacteria. This may explain why uptake-independent killing of macrophages was not reported in a previous study where cytochalasin D-treated macrophages were exposed to non-aggregated bacterial suspensions (26).

We observed that the plasma membrane of cytochalasin D-treated macrophages extends around *Mtb* aggregates and establishes a stable interaction with them. At this interface, the host-cell plasma membrane stains with Annexin V, a marker of membrane perturbation. This staining is not an artifact of cytochalasin D treatment, because local Annexin V staining is also observed in the small number of untreated macrophages that establish long-term contact with extracellular aggregates without internalizing them. We speculate that bacterial factors involved in triggering host-cell death may be concentrated over a smaller area of host-cell membrane when the bacteria are aggregated rather than dispersed. This may explain why aggregates are more efficient at inducing contact-dependent macrophage death compared to similar numbers of individual (non-aggregated) bacteria. Alternatively, aggregated bacteria might produce higher amounts of bacterial factors required to induce host-cell death or they might retain them more easily on their surface.

Previous studies have reported that the *Mtb* ESX-1 type VII secretion system and the ESX-1-secreted proteins EsxA/EsxB are required for escape from the phagosome and induction of host-cell death by intracellular bacteria (42, 5). Here, we show that EsxA/EsxB *expression* is also required to induce uptake-independent killing of macrophages upon contact with extracellular bacterial aggregates. Unexpectedly, however, we found that elimination of EsxA/EsxB *secretion* by deletion of *espA* (51, 52) has little effect on macrophage killing unless the ESX-1-secreted EspB protein is absent. We also found that EsxA/EsxB *expression* is required for secretion of the cleaved 50-kDa isoform but not the full-length 60-kDa isoform of EspB, in agreement with a recent study (56). Cleavage of EspB by MycP1 is required for full *Mtb* virulence (57), interaction of EspB with phospholipids (52), and oligomerization of EspB into channel-shaped heptamers (58, 59). These observations suggest that EspB can induce uptake-independent killing of macrophages in the absence of EsxA/EsxB *secretion* but not in the absence of EsxA/EsxB *expression*, which appears to be required for MycP1-dependent cleavage of EspB.

EspB secretion has been linked to *Mtb* virulence in a previous study comparing *Mtb* ESX-1 mutants with different secretion patterns, although the behavior of an isogenic *espB* deletion strain was not reported (57, 52). It has been suggested that EspB heptamers have been suggested to mediate secretion of other proteins (60, 58, 59, 61); however, this hypothesis is inconsistent with our proteomic analysis of the *espB* mutant’s secretome showing that deletion of *espB* does not affect the secretion of any other protein. Alternative models suggest that EspB heptamers may form membrane-spanning pores after contacting host-cell membranes, suggesting a possible mechanism for EspB-mediated cell death (61). However, addition of purified EspB to macrophages does not cause cell death (52), consistent with our observation that direct contact between *Mtb* aggregates and macrophages is required for killing and bystander macrophages are not affected even when they are in close proximity to *Mtb* aggregates. Similarly, while it is still debated whether purified EsxA alone has pore-forming activity on membranes (62–64), direct physical interaction between EsxA-producing strains and macrophages is also required to induce uptake-independent killing of macrophages, suggesting that secreted factors *per se* may not be sufficient to cause cell death.

A possible explanation for the contact-dependency of killing may be provided by our observation that PDIM, a complex lipid associated to the outer membrane of *Mtb* (65), is also required for killing of macrophages by *Mtb* aggregates, although it is not sufficient in the absence of ESX-1. Importantly, we found that the essential role of PDIM in contact-dependent killing is not linked to any detectable change in the *Mtb* secretome in the absence of PDIM. Previous reports have shown that PDIM can insert into host-cell membranes and modify their biophysical properties (66–68). PDIM has also been shown to increase the membranolytic activity of EsxA against liposomes (25, 63), but its interaction with EspB has not been investigated to the best of our knowledge. Association of PDIM with the bacterial surface may explain why direct physical contact between bacteria and host cells is required to induce macrophage death.

Contact with extracellular *Mtb* aggregates triggers local plasma membrane perturbation and cytoplasmic calcium accumulation in macrophages followed by cell death. We observed that cells that display perturbed membrane foci at the site of contact with *Mtb* aggregates progressively accumulate cytoplasmic calcium before death. PDIM-deficient bacteria cause local plasma membrane perturbation without causing cell death, suggesting that membrane perturbation *per se* is not sufficient to induce cell death. Membrane perturbation in macrophages requires secretion of EsxA/EsxB or EspB and may be required for cytoplasmic calcium accumulation because mutations that eliminate membrane perturbation (*esxA* and *espA espB* mutants) also eliminate calcium accumulation. Conversely, calcium accumulation seems not to be required for membrane perturbation because loss of PDIM eliminates calcium accumulation without affecting membrane perturbation. Taken together, these results suggest that although PDIM and secreted EsxA/EsxB and EspB all participate in uptake-independent killing of macrophages after contact with extracellular *Mtb* aggregates, their roles are somewhat different.

We propose two alternative models that could explain these observations. EsxA/EsxB- or EspB-dependent plasma membrane perturbation may facilitate PDIM insertion into the membrane, which could potentially affect membrane permeability to ions such as calcium. Alternatively, EsxA (25) or EspB may require PDIM to potentiate their membranolytic activities and to alter membrane permeability to ions. Both PDIM and a fraction of secreted EsxA and EspB are physically associated with the bacterial cell surface (69–71, 56), which may explain why direct contact of *Mtb* with macrophages is required for induction of host-cell death while bystander macrophages are not affected.

A general caveat of models based on analysis of *esxA* or *espA* mutants is that EsxA and EspA are mutually dependent for their secretion and it is thus unclear whether some of the phenotypes observed may depend on one or the other protein. EspA has been shown to have an essential role in EsxA secretion (51). However, in contrast to EsxA (42), we are not aware of any published evidence that EspA interacts with host-cell membranes and exerts membranolytic activity. For these reasons, it is generally assumed that EsxA is the secreted bacterial effector most responsible for damaging host-cell membranes (5, 72, 19).

Uptake-independent death of macrophages by extracellular *Mtb* aggregates can be partially suppressed by treating the infected cells with pyroptosis inhibitors targeting NLRP3 or Caspase 1. Inhibition of this pathway reduces cell death without affecting either formation of Annexin V-positive membrane domains or intracellular calcium accumulation. These results suggest that extracellular *Mtb* aggregates induce imbalance of cytoplasmic calcium or other ions in contacted macrophages, leading to inflammasome activation and pyroptotic cell death. Intracellular ion imbalances and pyroptosis have similarly been observed in macrophages after plasma membrane damage caused by intracellular *Mtb* (30).

Our work highlights the importance of *Mtb* aggregation during host cell infection and demonstrates how *Mtb* aggregates can evade phagocytosis by inducing contact-dependent but uptake-independent death in macrophages. *Mtb* aggregation has been shown to play an important role in pathogenesis *in vivo* (73, 74), and several groups have shown that macrophages that internalize single bacteria are able to survive for days whereas uptake of *Mtb* aggregates typically results in rapid death (19, 20). Although *Mtb* infections can start with very few individual bacteria (2), these bacteria can form aggregates when they grow inside host cells (9, 18) or on the debris of dead host cells (8). Presence of intracellular and extracellular *Mtb* aggregates have been documented in live and necrotic cells in the lungs of mouse, rabbit and guinea pig models of infection already within the first month post infection (9–11, 74). Moreover, aggregates have also been observed in human lung lesions and sputum samples (12–18), confirming their potential relevance in human tuberculosis infection.

We propose that uptake-independent induction of macrophage death by *Mtb* aggregates may promote the propagation of *Mtb* at different stages of infection. At early stages, release of *Mtb* aggregates upon death of infected naïve macrophages (killing “from the inside”) may allow rapid replication of extracellular bacteria on the host-cell debris. Subsequently, uptake-independent killing “from the outside” may allow these extracellular aggregates to evade phagocytosis by less-permissive macrophages as host immunity begins to develop (75, 76). Even when *Mtb* aggregates are successfully internalized by macrophages, they may still evade intracellular host defenses by inducing rapid host-cell death “from the inside” (20). At later stages, toxicity of extracellular *Mtb* aggregates may also participate in the necrotic processes required for expansion of lesions, formation of large extracellular bacterial pellicle in open necrotic cavities and bacterial spillover into the airways (9, 12, 14). *Mtb* aggregates that spill into the airways may induce death of newly recruited macrophages, thereby contributing to evasion of host immunity, exhalation of live bacteria, and transmission of the infection (77). Based on our results with BTP15, a small-molecule inhibitor of ESX-1 (53), we propose that bacterial spreading within the lung could be suppressed by novel therapies targeting virulence factors, such as ESX-1 or PDIM, that are required for uptake-independent killing of host macrophages by extracellular *Mtb* aggregates. By reducing host-cell death, such therapies could also suppress the formation of necrotic lesions, where large numbers of extracellular bacteria may grow rapidly on the debris of dead host cells (8) within an environment that is poorly penetrated by antibiotics (9, 78, 79).

## Materials and methods

### Bacterial strains and growth conditions

Strains used in this study include WT *Mtb* H37Rv Pasteur, WT *Mtb* Erdman, various mutants, complemented and fluorescent strains listed in Table S1. All strains were cultured in liquid Middlebrook 7H9 liquid medium (Difco) supplemented with 10% ADC (Difco), 0.5% glycerol and 0.02% Tyloxapol at 37 °C with shaking. As solid medium Middlebrook 7H10 (Difco) supplemented with 10% OADC enrichment (Becton Dickinson) and 0.5% glycerol was used. When required, kanamycin, hygromycin, zeocin were added to the medium at a final concentration of 25 μg/ml, 50 μg/ml and 25 μg/ml, respectively.

### Generation of fluorescent, mutant and complemented strains

All the tdTomato fluorescent strains were developed by electroporating the parental strains with the integrative pND257 plasmid. pND257 is a mycobacterial L5-based integrative plasmid in which the gene encoding TdTomato was cloned downstream of a strong constitutive promoter and confers resistance to kanamycin.

Transformants were selected plating on 7H10 plates + kanamycin and verified by fluorescence microscopy.

The *espB* mutants were constructed using the ORBIT strategy (80). The parental strains *Mtb* Erdman wild-type and *espA*-tn were electroporated with the plasmid pKM461 (Addgene, #108320) expressing tetracycline-inducible Che9c phage RecT annealase and Bxb1 phage integrase (80). Transformants were selected plating on 7H10 plates + kanamycin. Cells containing pKM461 were grown in liquid medium + kanamycin and supplemented with 500 ng/ml anhydrotetracycline when the culture reached an OD_600_ of 0.3. After 24 h induction, bacteria were electroporated with the payload plasmid pKM496 (Addgene, #109301) (80) and 1 μg of the Ultramer DNA oligonucleotide (IDT) targeting *espB* (agcgtcaacgcccgggcgacatgcgggtccaattcgtccatgctcacttcgactccttactgtcctggcgGGTTTGTACCGTACACCACTGAGACCGCGGTGGTTGACCAGACAAACCctgcgactgcgtcatatcggatcatcctccttagtgctatagccattatcgtcgctaaactgaaaggttc). Recombinants were selected plating on 7H10 plates + zeocin. The recombination was verified by PCR analysis. For the selected clones the plasmid pKM461 was cured by successive growth in kanamycin-free medium. Loss of plasmid was verified by inoculating bacteria in liquid medium + kanamycin.

For complementation of the *espB* mutant strains the pMV261_*espB* plasmid was developed. A DNA region covering the *espB* gene plus 250 bp upstream of the start codon was amplified by PCR from *Mtb* Erdman genomic DNA using the primers espBcompF and espBcompR (Table S2). The resulting PCR product was integrated in the pMV261 (81) plasmid digested with AatII and EcoRI using the Gibson Assembly strategy according to the manufacturer protocol (NEB).

### Cell cultures

BMDMs were differentiated from cryopreserved bone marrow stocks extracted from femurs of 6- to 8-week-old C57BL/6 and mTmG mice (Jackson Laboratory). The bone marrow was cultured in petri dishes in BMDM differentiation medium (DMEM with 10% FBS, 1% sodium-pyruvate, 1% GlutaMax and 20% L929-cell-conditioned medium as a source of granulocyte/macrophage colony stimulating factor). After 7 days, adherent cells were gently lifted from the plate using a cell scraper, resuspended in BMDM culture medium (DMEM with 5% FBS, 1% sodium-pyruvate, 1% GlutaMax and 5% L929-cell-conditioned medium) and seeded in a 35 mm 4-compartment Ibidi μ-dishes or in Ibidi 24 well-plates and incubated at 37°C, 5% CO_2_ before further experiments.

### Macrophages infection

For infection with single non-aggregated bacteria, 1 ml of *Mtb* culture at OD_600_ 0.4-0.8 was pelleted, resuspended in 200 μl of macrophage medium and passed through a 5 μm filter to eliminate bacterial aggregates. The resulting single-cell suspension was used to infect BMDMs macrophages at an MOI of 1:1. After 4 hours of infection, macrophages were washed extensively with macrophage medium to remove extracellular bacteria.

For infection with bacterial aggregates, 1 ml of *Mtb* culture at OD_600_ 0.4-0.8 was pelleted and resuspended in 1 ml of macrophage medium. A 1:100 dilution of this solution was used to infect macrophages.

For spinfection experiments, a single-cell suspension of *Mtb* or a solution of bacterial aggregates were added to BMDMs seeded in 24 well-plates. After centrifugation at 500 g for 5 minutes, the medium was removed and replaced with fresh medium. When required, 4 μM cytochalasin D (Invitrogen), 1:1000 diluted Annexin V-FITC (BioLegend) or Annexin V-647 (BioLegend), 1:1000 Draq7 (BioLegend), 50 μM BTP15 (gift from Prof. S. Cole), 50 μM Z-IETD-FMK (Selleck), 5 μM GSK872 (Abcam), 10 μM MCC950 (Selleck), 50 μM VX765 (Abcam), 10 μM dantrolene (Abcam), 10 mM CaCl_2_, 5 μM BAPTA-AM (Selleckchem), 20 ng/ml TNFα (R&D systems),10 μg/ml cycloheximide (Sigma-Aldrich), 50 ng/ml LPS (Sigma-Aldrich), 0.5 mM ATP (Sigma-Aldrich), 5 μM z-VAD-FMK (InvivoGen), 20 50 nM SM164 (Selleckchem) were added at the time of infection and kept in the medium of the macrophages during the course of the experiment. When macrophages were stained with Oregon Green 488 Bapta-1 AM (ThermoFisher Scientific), the staining was performed adding 5 μM dye in PBS to the cells and incubating for 30 min at 37°C, 5% CO_2_. Macrophages were then washed three time in PBS to remove any residual extracellular dye before infection.

### Immunofluorescence on macrophages infected with *Mtb*

Macrophages were fixed for 2 h with 4% paraformaldehyde in PBS at 24 hours post infection. Cells were then washed three times with PBS and permeabilized for 15 minutes with 2% BSA, 2% saponine, 0.1% Triton x-100 in PBS at RT. After three washes with PBS, samples were blocked for 1 hour with 2% BSA in PBS at RT. Rabbit primary antibodies anti-cleaved Caspase-8 (Cell Signaling), ASC/TMS1 (Cell Signaling), cleaved Caspase-1 (Cell Signaling), phospho-RIP3 (Cell Signaling), phospho-MLKL (Cell Signaling), mouse CD45.1-AlexaFluor647 (BioLegend) were diluted 1:300 in 2% BSA in PBS and used for primary staining of the cells overnight at 4°C. After three washes in 2% BSA in PBS, samples (excluded those stained with anti-mouse CD45.1-AlexaFluor647) were incubated with a secondary goat anti-rabbit antibody conjugated to Alexa Fluor 647 (ThermoFisher Scientific) diluted 1:300 in 2% BSA in PBS for 1 hour at RT. Samples were stained with 1:1000 Hoechst (ThermoFisher Scientific) and stored in PBS at 4°C before imaging by fluorescence microscopy.

### Microscopy on macrophages infected with *Mtb*

For experiments quantifying cell viability, bacterial growth and Annexin V staining, infected BMDMs were imaged by time-lapse microscopy for up to 132 hours at 1 or 2-hour intervals with a 20x air objective on a DeltaVision microscope or a 40x air objective on a Nikon Ti2 microscope. 3 x 1.5 μm z-stacks were acquired on multiple XY fields.

For experiments monitoring the fluorescence of the infected cells (Annexin V and Draq7 staining, Oregon Green 488 Bapta-1 AM fluorescence, mTmG BMDMs’ membrane fluorescence), infected BMDMs were imaged by time-lapse microscopy for up to 24 hours at 20-minute intervals with a 63x oil-objective on a DeltaVision microscope. 5 x 1 μm z-stacks were acquired on multiple XY fields. Our DeltaVision microscope is equipped with FITC (Ex 490/20, Em 525/36), TRITC (Ex 555/25, Em 605/52), Cy-5 (Ex 555/25, Em 605/52) and DAPI (Ex 438/24, Em 475/24) dichroic filters. On the Nikon Ti2 microscope GFP (Em 480/30, Ex 535/45) and mCherry (Em 560/40, Ex 635/60) dichroic filters were used. A stage-top incubator (Okolab) was used to maintain the cells at 37°C in a humidified environment. Air mixed to 5% CO_2_ was supplied using an Okolab gas mixer. One half of the macrophage medium was refreshed every 3 days through custom tubing connected to the lid of the Ibidi μ-dish. Immunostained samples were imaged with a 63x oil-objective on a DeltaVision microscope. 5 x 1 μm z-stacks were acquired on multiple XY fields.

Confocal images on fixed samples were acquired using a Leica SP8 microscope. Top and bottom z-coordinates for all the selected cells were assigned manually and z-stacks were acquired every 0.29 μm.

### Image analysis

The microscopy images and time-series were analyzed using the FIJI software from the ImageJ package (82). The z-stacks acquired for the individual fluorescence channels were projected into one image using a maximum intensity projection. A manual threshold was set to segment the bacteria. The area above the threshold was measured and used as a proxy for the number of bacteria for each time point. To quantify the bacterial growth rate, an exponential curve was fitted to the data. In experiment without cytochalasin D treatment, bacteria were considered intracellular when they overlap with a macrophage in the bright-field channel, otherwise they were annotated as extracellular. Macrophage infection status, Annexin V staining and viability were monitored by visual analysis of the time-lapse image series.

Regions of interest corresponding to individual macrophages were manually drawn onto the bright-field images and transferred to fluorescence images. Macrophages were considered infected when the region of interest overlaps with a fluorescent region in the channel used to image bacteria. Similarly, macrophages were considered Annexin V-positive upon appearance of a positive regions in the corresponding fluorescence channel. Time of death corresponds to the frame where a macrophage stops moving and lyses. The time interval between initial infection and the appearance of the Annexin V-positive domain or the cell death was calculated as the difference between the first frame where a macrophage was infected and the frame where the host cell became Annexin V-positive or died. For experiments with Oregon Green 488 Bapta-1 AM, the z-stacks acquired in the green channel were projected into one image using a “sum slices” projection. A background subtraction was performed by subtracting from the fluorescence images a copy of the same images on which a Gaussian blur of 100 μm radius had been applied. Regions of interest (ROIs) corresponding to individual macrophages were manually drawn onto the phase images and transferred to the fluorescence images. The Oregon Green 488 Bapta-1 AM fluorescence of each macrophage was quantified for each frame as average fluorescence intensity and normalized to the first frame imaged (for bystander cells) or to the frame of infection (for infected cells). For images acquired from cells stained with anti-ASC or anti-cleaved Caspase8 (cCasp1) antibodies, ASC positive or cCasp8 positive cells were counted through manual analysis of the fluorescence images. For images acquired from cells stained with anti-cleaved Caspase1, anti-phosphorylated RIP3 or anti-phosphorylated MLKL antibodies, the z-stacks acquired in the Cy5 channel were projected into one image using a “sum slices” projection. ROIs corresponding to individual macrophages were drawn onto the phase images and transferred to the fluorescence channel to measure the background subtracted median fluorescence for each cell. Cells whose ROIs overlap with bacteria in the bacterial fluorescence channel were annotated as infected.

Confocal images were processed with ImageJ for 3D reconstruction. A Gaussian Blur filter with a 2 pixel radius was applied to the channel corresponding to the macrophage membrane. The TransformJ:Rotate plugin was used to resample isotropically all the channels using a Quantic B-Spline interpolation. The 3D visualization was obtained using the 3D Script – Interactive Animation tool and a combined transparency rendering.

### Scanning electron microscopy (SEM)

Cells were fixed for 2 hours with 1.25% glutaraldehyde in 0.1 M phosphate buffer, pH 7.4, and then washed in cacodylate buffer before a secondary fix for 30 minutes in 0.2 % osmium tetroxide in the same buffer. The cells were then dehydrated in a graded alcohol series and dried by passing them through the supercritical point of carbon dioxide in a critical point dryer (Leica Microsystems CPD300). They were then coated with a 2nm layer of osmium metal using an osmium plasma coater (Filgen OPC60). Images of the cells were taken with a field emission scanning electron microscope (Merlin, Zeiss NTS).

### Colony Forming Units (CFUs) counting

For CFUs enumeration over time, macrophages in individual wells were infected at time point zero. At the different time points, the medium of the infected cells was collected for plating. The macrophages were lysed with 0.5% TritonX-100 in PBS for 5 minutes at room temperature, and the lysate was collected and mixed to the medium. Serial dilutions of this mixture were plated on solid medium and incubated at 37 °C. After three weeks CFUs were counted.

### RNA extraction and Quantitative Real-Time PCR (qRT-PCR)

10 ml Mtb cultures in mid-logarithmic phase were pelleted by centrifugation, resuspended in 800 μl of TRIzol Reagent (ThermoFisher) and added to a 2-ml screw-cap tube containing zirconia beads (BioSpec Products). Cells were disrupted by bead-beating three times for 30 seconds. The cell lysate was transferred to a new tube, mixed with 200 μl of chloroform and spun down for 15 min at 12,000 g. The top aqueous phase was collected and mixed with an iso-volume of isopropanol. The RNA was precipitated by centrifugation for 10 min at 15,000 g, washed with 75% ethanol, air-dried and resuspended in DEPC-treated water. DNase treatment was carried out using TURBO DNA-free kit (Invitrogen) according to the manufacturer’s protocol. The RNA was transcribed to cDNA using the SuperScript IV First-strand Synthesis System (ThermoFisher) according to the manufacturer’s recommendations. qRT-PCR were performed on an ABI 7900HT instrument, using SYBRGreen PCR Master Mix (Applied Biosystems), according to the manufacturer’s instructions. Relative mRNA levels were calculated using the ΔΔCt method, normalizing transcripts levels to *sigA* signals. The sequences of the primers used are listed in table S2.

### Protein extracts and immunoblotting

*Mtb* was inoculated in 10 ml cultures (or 30 ml cultures for proteomics samples) in Sauton’s medium supplemented with 0.05% Tween 80 at a starting OD_600_ of 0.1 and incubated at 37°C with shaking. After 4 days the cells were harvested by centrifugation, washed with PBS and resuspended in 10 ml (or 30 ml cultures for proteomics samples) of Sauton’s medium without Tween 80 and incubated at 37°C with agitation for 4 days. Cultures were pelleted by centrifugation and the collected culture media were filtered through a 0.2-μm-pore-size filter. Culture filtrates (secretomes) were concentrated 50x using Amicon Ultra centrifugal filters with 3-kDa cut-off (Millipore) respectively. To prepare total “bacterial lysates”, the bacterial pellets were resuspended in TBS (20 mM Tris + 150 mM NaCl) plus cOmplete mini EDTA free protease inhibitors (Roche) and disrupted by bead beating four times for 30 seconds with zirconia beads (BioSpec Products). After clarification by centrifugation, the samples were filtered with a 0.2-μm-pore-size filter. Total protein concentration in all the samples was determined using a Pierce BCA assays (ThermoFisher) with bovine serum albumin as the standard. Protein samples were resolved in NuPAGE 4 to 12% bis-tris gels (Invitrogen) under reducing conditions. Proteins were transferred from the gel to a nitrocellulose membranes using the iBlot Dry Blotting System (Invitrogen) according to the manifacturer’s instructions.

Membranes were blocked for 1 h at room temperature with TBS + 3% BSA fraction V and then incubated overnight with the primary antibody diluted in TNT (TBS + 0.1% Tween-20) + 2% BSA fraction V at 4°C. Membranes were washed with TNT, incubated with the secondary antibody in TNT+ 2% BSA for 30 min at room temperature, washed with TNT, and developed.

### Mass spectrometry on protein samples

Mass spectrometry-based proteomics-related experiments were performed by the Proteomics Core Facility at EPFL. Each sample was digested by filter aided sample preparation (FASP) (83) with minor modifications. Proteins (40 μg) were reduced with 10 mM TCEP in 8 M Urea, 0.1 M Tris-HCl pH 8.0 at 37°C for 60 min and further alkylated in 40 mM iodoacetamide at 37°C for 45 min in the dark. Proteins were digested overnight at 37°C using 1/50 w/w enzyme-to-protein ratio of mass spectrometry grade trypsin gold and LysC. Generated peptides were desalted in StageTips using 6 disks from an Empore C18 (3 M) filter based on the standard protocol (84). Purified peptides were dried down by vacuum centrifugation.

Samples were resuspended in 2% acetonitrile (Biosolve), 0.1% FA and nano-flow separations were performed on a Dionex Ultimate 3000 RSLC nano UPLC system (Thermo Fischer Scientific) on-line connected with an Orbitrap Exploris Mass Spectrometer (Thermo Fischer Scientific). A capillary precolumn (Acclaim Pepmap C18, 3 μm-100Å, 2 cm x 75μm ID) was used for sample trapping and cleaning. A 50cm long capillary column (75 μm ID; in-house packed using ReproSil-Pur C18-AQ 1.9 μm silica beads; Dr. Maisch) was then used for analytical separations at 250 nl/min over 150 min biphasic gradients. Acquisitions were performed through Top Speed Data-Dependent acquisition mode using a cycle time of 2 seconds. First MS scans were acquired with a resolution of 60’000 (at 200 m/z) and the most intense parent ions were selected and fragmented by High energy Collision Dissociation (HCD) with a Normalized Collision Energy (NCE) of 30% using an isolation window of 2 m/z. Fragmented ions were acquired with a resolution 15’000 (at 200m/z) and selected ions were then excluded for the following 20 s.

Raw data were processed using MaxQuant 1.6.10.43 (85) against the Uniprot Reference Proteome Mycobacterium Tuberculosis Erdman strain (4222 sequences, LM201030). Carbamidomethylation was set as fixed modification, whereas oxidation (M), phosphorylation (S, T, Y), acetylation (Protein N-term) and glutamine to pyroglutamate were considered as variable modifications. A maximum of two missed cleavages were allowed and “Match between runs” option was enabled. A minimum of 2 peptides was required for protein identification and the false discovery rate (FDR) cutoff was set to 0.01 for both peptides and proteins. Label-free quantification and normalisation was performed by Maxquant using the MaxLFQ algorithm, with the standard settings (86). The statistical analysis was performed using Perseus version 1.6.12.0 (87) from the MaxQuant tool suite. Reverse proteins, potential contaminants and proteins only identified by sites were filtered out. Protein groups containing at least 2 valid values in at least one condition were conserved for further analysis. Missing values were imputed with random numbers from a normal distribution (Width: 0.3 and Down shift: 1.8 sd). A two-sample t-test with permutation-based FDR statistics (250 permutations, FDR=0.05, S0=1) was performed to determine significant differentially abundant candidates. The mass spectrometry proteomics data have been deposited to the ProteomeXchange Consortium (88) via the PRIDE (89) partner repository Data relative to the *Mtb* Erdman strains have the dataset identifier PXD035080 (username, password: 0BdFQzrQ). Data relative to the *Mtb* H37Rv strains have the dataset identifier PXD035082 (username, password: CMqe5spP).

### PDIM extraction

Phthiocerol dimycocerosates (PDIM) synthesis was evaluated by thin-layer chromatography (TLC) of bacterial lipids (90). 10 ml of exponential-phase bacteria were labeled with 2 μCi of [14C]-propionate (Campro Scientific) for 48 hours. Lipids were extracted from the pellets of the radiolabelled culture using 5 ml of methanol : 0.3% NaCl (10:1) and 5 ml of petroleum ether. The mixture was vortexed for 3 min, centrifuged at 930 g for 10 minutes, and the upper layer was collected. A second extraction on the lower phase was carried out with 5 ml of petroleum ether. The combined extracts were inactivated with an iso-volume of chloroform for 1 hour and concentrated through evaporation. The samples were spotted on a 5 x 10 cm TLC silica gel 60 F254 (Merck) and developed in petroleum ether:diethyl ether (9:1). The developed TLC plate was exposed to an Amersham Hyperfilm ECL (GE Healthcare) for chemioluminescence analysis and visualized with a Typhoon Scanner (GE Lifesciences).

## Acknowledgments

This work was supported by grants to J.D.M. from the Swiss National Science Foundation (310030B_176397). This work has received support from the Innovative Medicines Initiatives 2 Joint Undertaking grant No 853989 “ERA4TB” (https://era4tb.org/). C.T. was supported by funding from the European Union’s Horizon 2020 research and innovation program under the Marie Skłodowska-Curie grant agreement No. 665667. VIDO receives operational funding from the Government of Saskatchewan through Innovation Saskatchewan and the Ministry of Agriculture and from the Canada Foundation for Innovation through the Major Science Initiatives for its CL3 facility.

We would like to thank Dr. Maria Pavlou, Dr. Florence Armand and Jonathan Pittet from the Proteomics Core Facilities at the School of Life Sciences of EPFL for their help in processing and analyzing the *Mtb* secretome samples. We acknowledge Anaëlle Dubois, Stéphanie Rosset, Marie Croisier and Prof. Graham W. Knott from the EPFL Biological Electron Microscopy Facility for their help in preparing and imaging samples by SEM. We acknowledge Dr. Nicholas Chiaruttini, José Artacho and Dr. Arne Seitz from the EPFL Bioimaging and Optics Platform for their help with confocal microscopy and image analysis. We thank Dr. Daniel Sage for help with image analysis; Dr. Catherine Astarie-Dequeker for sharing the H37Rv Δ*esxA* strains; Prof. Stewart Cole and Prof. Jeffrey Chen for providing antibodies, *Mtb* strains, and BTP15. Finally, we thank all the members of the McKinney group for fruitful discussions and feedback on the manuscript.

## Author Contributions

**Chiara Toniolo:** Conceptualization, Investigation, Data curation, Writing - original draft, Writing - review & editing, Visualization.

**Neeraj Dhar**: Conceptualization, Writing - review & editing, Supervision.

**John D McKinney:** Conceptualization, Writing - review & editing, Supervision, Funding acquisition.

The authors declare that they have no conflict of interest.

## Supplementary Information

**Supplementary Movie S1. Time-lapse microscopy on macrophages infected with *Mtb*.**

BMDMs were infected with *Mtb* expressing tdTomato (magenta) and imaged by time-lapse microscopy at 1-hour intervals for 146 hours. After uptake, single bacteria grow inside the infected macrophage and eventually induce death and lysis. Extracellular *Mtb* aggregates continue growing on the debris of the dead host cells and attract bystander macrophages. Scale bar, 40 μm.

**Supplementary Movie S2. Representative example of a *Mtb*-infected macrophage that dies.**

BMDMs infected with individual *Mtb* bacteria (magenta) were imaged by time-lapse microscopy at 20 minute-intervals for 144 hours. Scale bar, 10 μm. Example of a macrophage that survives for 54h20 before dying. Dying macrophages typically stop moving, shrink, lose membrane integrity and stain positive for the membrane impermeable dye Draq7 (blue). When macrophages are alive, the individual intracellular bacteria move over time following the physiological movements of the host cells. Upon macrophage death, bacteria stop moving and grow on the debris of the dead cell.

**Supplementary Movie S3. Time-lapse microscopy on macrophages infected with *Mtb* aggregates from axenic culture.**

BMDMs were infected with aggregates of *Mtb* Erd-tdTomato (magenta) from axenic cultures and imaged by time-lapse microscopy at 1-hour intervals for 60 hours. Scale bar, 40 μm.

**Supplementary Movie S4. Representative example of a macrophage that die upon interaction with a *Mtb* aggregate that undergoes fragmentation.**

BMDMs infected with aggregates of *Mtb* (magenta) were imaged by time-lapse microscopy at 20 minute-intervals for 20 hours. Scale bar, 10 μm. Example of a BMDM that interacts with a *Mtb* aggregate (magenta), fragments it and redistributes the bacteria. The macrophage dies at 17h40 post interaction with the aggregate, when it stops moving, loses membrane integrity and stains positive for the membrane impermeable dye Draq7 (blue).

**Supplementary Movie S5. Representative example of a macrophage that die upon interaction with a *Mtb* aggregate that does not undergo fragmentation.**

BMDMs infected with aggregates of *Mtb* (magenta) were imaged by time-lapse microscopy at 20 minute-intervals for 20 hours. Scale bar, 10 μm. Example of a BMDM that interacts with a *Mtb* aggregate (magenta). The bacterial aggregate folds and changes shape during the interaction with the macrophages but it does not undergo fragmentation. The macrophage dies at 4h40 post interaction with the aggregate, when it stops moving, loses membrane integrity and stains positive for the membrane impermeable dye Draq7 (blue).

**Supplementary Movie S6. Time-lapse microscopy on cytochalasin D-treated macrophages infected with aggregates of *Mtb*.**

Cytochalasin D-treated BMDMs were infected with aggregates of *Mtb* Erd-tdTomato (magenta) and imaged by time-lapse microscopy at 1-hour intervals for 60 hours. Scale bar, 40 μm.

**Supplementary Movie S7. Representative example of a cytochalasin D-treated macrophage that die upon contact with an extracellular *Mtb* aggregate.**

**(Left panel)** Cytochalasin D-treated BMDMs establish stable contact with extracellular *Mtb* aggregates (magenta). The bottom macrophage dies at 19h40 post interaction with the aggregate, when it stops moving, loses membrane integrity and stains positive for the membrane impermeable dye Draq7 (blue). **(right panel)** AnnexinV-positive domains (yellow) appear at the site of contact with the *Mtb* aggregates at 2h00 and 2h40 and stay bright until the end of the movie. When the bottom macrophage dies it loses membrane integrity and the whole cell stains positive for AnnexinV.

**Supplementary Movie S8. Representative example of a macrophage that shows multiple membrane perturbation events at the site of contact with an *Mtb* aggregate.**

Cytochalasin D-treated BMDMs establish stable contact with extracellular *Mtb* aggregate (magenta). The macrophage dies at 8h40 post interaction with the aggregate (Draq7-positive nucleus in blue in right panel). Multiple peaks in Annexin V fluorescence are observed at the site of contact with the *Mtb* aggregate (1^st^ at 1h00; 2^nd^ at 2h00; 3^rd^ at 7 h). Upon death of the macrophage, the *Mtb* aggregate retains some Annexin V-positive material on its surface and slowly decreases in fluorescence.

**Supplementary Figure S1.**
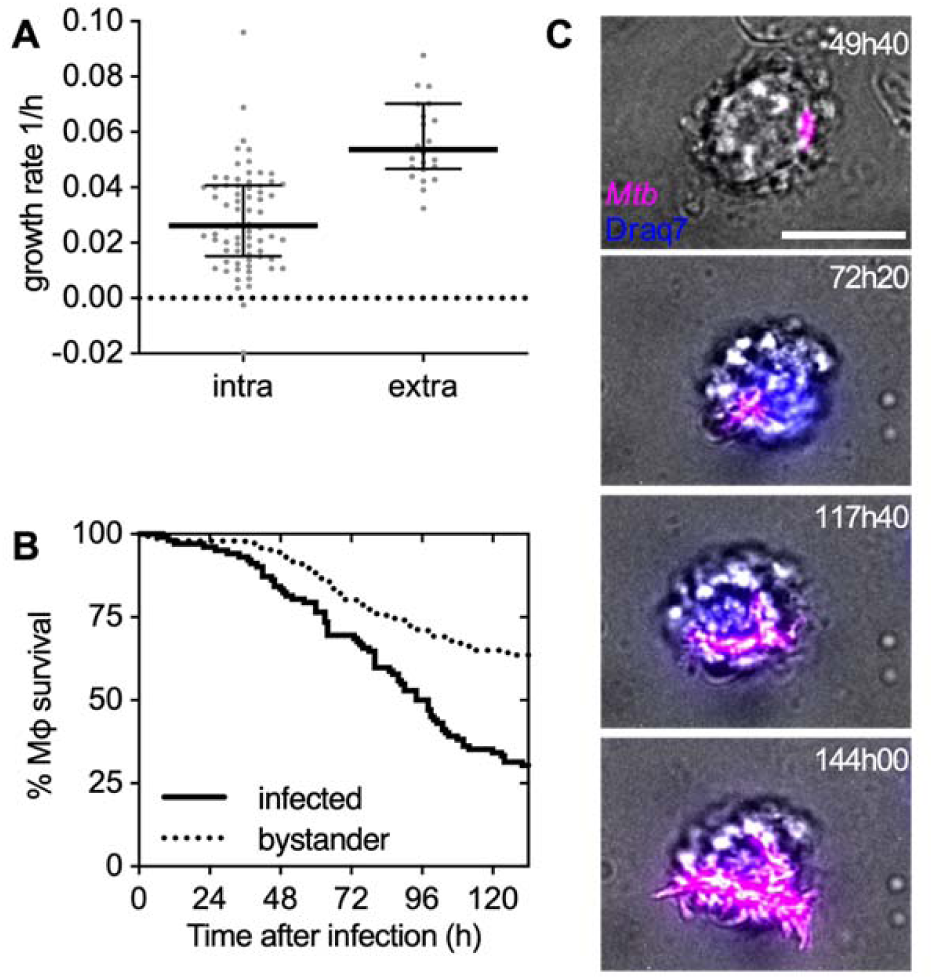
Intracellular growth of *Mtb* results in death and lysis of the infected macrophage, rapid extracellular growth on the host-cell debris, and formation of large extracellular *Mtb* aggregates. **(A-C)** BMDMs were infected with *Mtb* Erd-tdTomato and imaged by time-lapse microscopy at 1- or 2-hour intervals for at least 132 hours. **(A)** The bacterial growth rate was calculated by measuring the fluorescent area over time of individual *Mtb* microcolonies growing inside a macrophage (intra) or on the debris of a lysed macrophage (extra). Each symbol represents an *Mtb* microcolony. Black lines indicate median values and interquartile ranges. **(B)** Percentage survival over time of infected versus uninfected bystander macrophages. **(C)** Representative example of an intracellular *Mtb* microcolony that after lysis of the host macrophage (Draq7 positive cell) grows on the debris of the dead cell. Scale bar, 10 μm. Representative snapshots from movie S2.

**Supplementary Figure S2.**
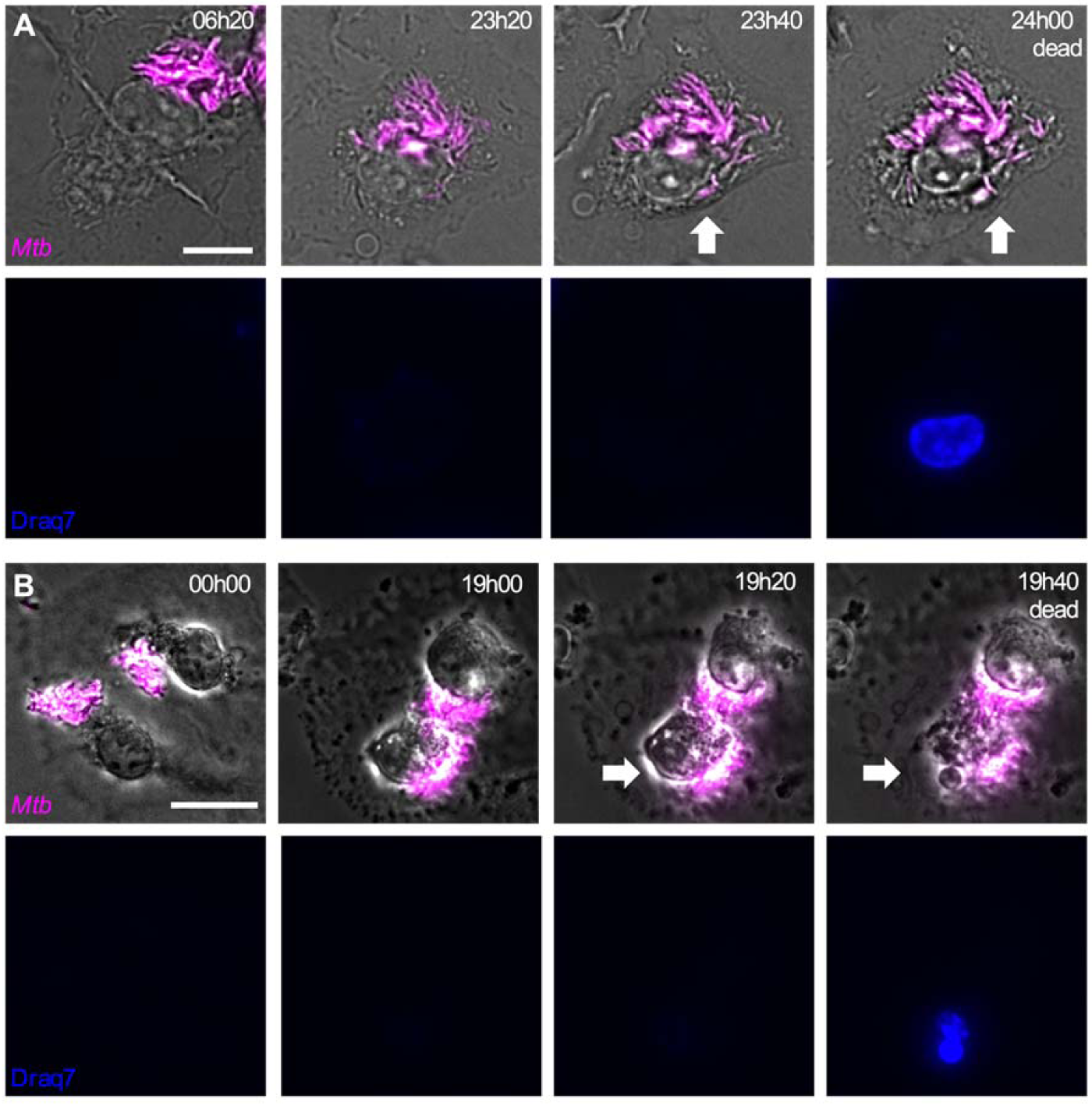
The macrophage time-of-death identified in brightfield images corresponds to the time-of-death defined by a fluorescent marker of cell death. Although it may be difficult to distinguish live and dead cells in individual snapshots, cell-death events are easily identifiable by comparing adjacent frames in image series obtained by time-lapse microscopy (supplementary movies S2, S7). When macrophages die, they rapidly shrink, lose membrane integrity (white arrows), and stop moving. In brightfield images we define the time-of-death for individual macrophages as the first image frame in which a cell stops moving and loses membrane integrity. At this time, also the intracellular bacteria identified in the fluorescent images and the intracellular structures (vesicles, nucleus) visible in the brightfield images stop moving. The macrophage time-of-death identified in brightfield images overlaps to the time-of-death defined by Draq7 (in blue), a cell-impermeable fluorescent marker commonly used to distinguish dead cells. **(A)** Representative example of a dying *Mtb*-infected macrophage. Snapshots from movie S4. **(B)** Representative example of cytochalasin D-treated macrophages that dies upon contact with an extracellular *Mtb* aggregate. Snapshots from movie S7. Scale bars, 10 μm.

**Supplementary Figure S3.**
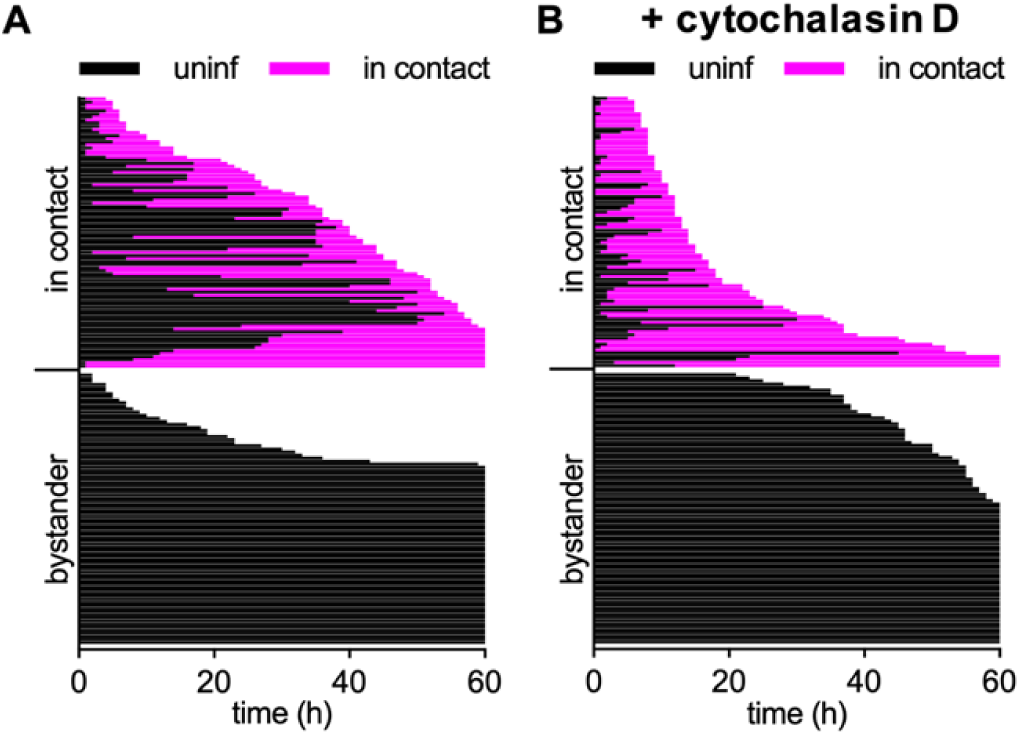
Physical interaction between bacterial aggregates and macrophages is required to induce death of the infected cells. Untreated **(A)** or cytochalasin D-treated **(B)** BMDMs infected with aggregates of *Mtb* Erd-tdTomato and imaged by time-lapse microscopy at 1-hour intervals for 60 hours. Each line represents the life span of an individual cell; the fraction of the line in black represents the time spent as uninfected, whereas the fraction of the line in magenta represents the time spent interacting with an *Mtb* aggregate. n=88 for each subclass (bystander, in contact).

**Supplementary Figure S4.**
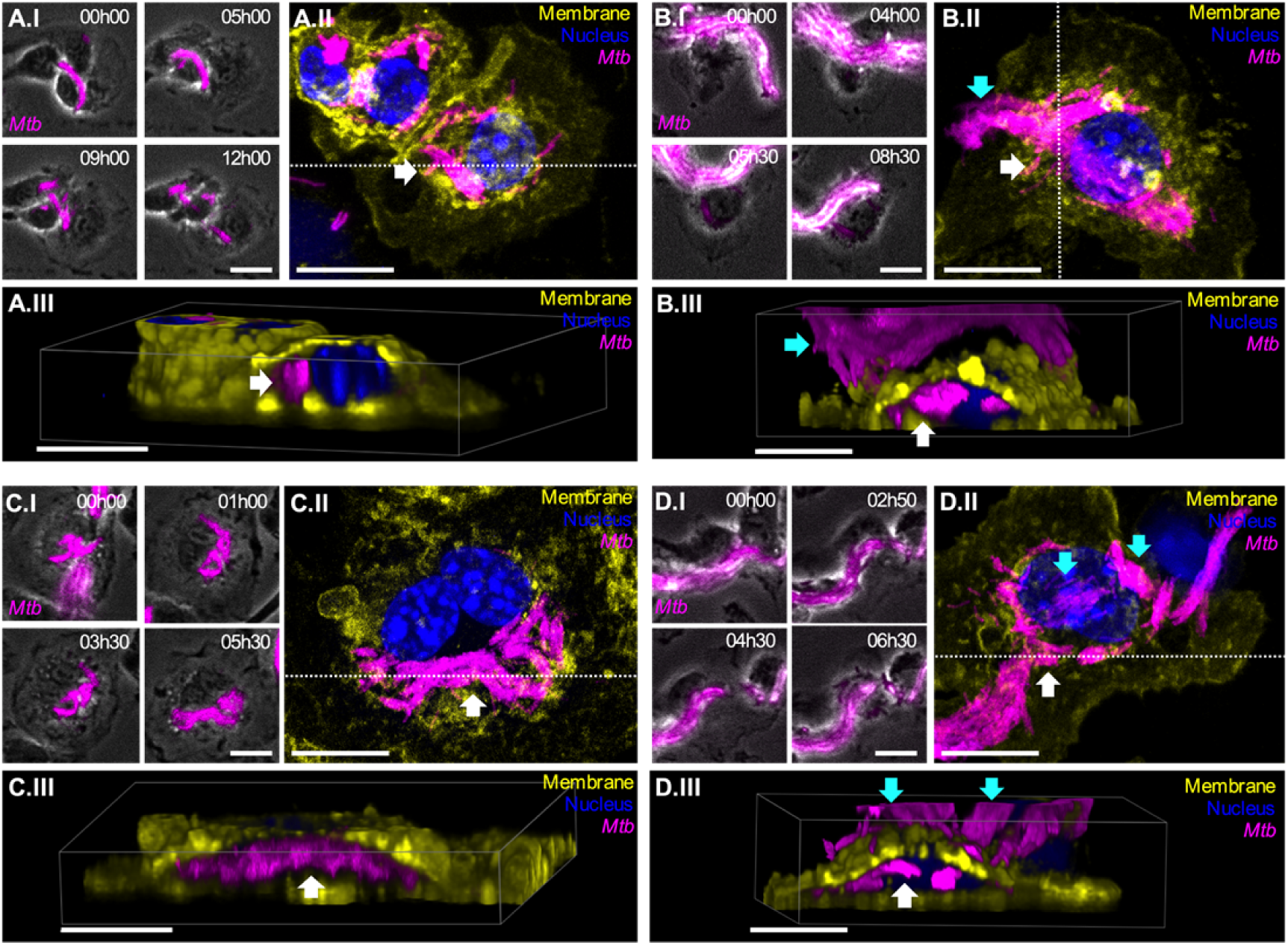
Correlative time-lapse microscopy and immunofluorescence on BMDMs infected with *Mtb* aggregates. **(A-D)** BMDMs infected with aggregates of *Mtb* were imaged by time-lapse microscopy (every 30 min for up to 13.5h) followed by fixation, immunostaining and imaging by confocal microscopy. *Mtb* in magenta, nuclei stained with Hoecsth (blue), membrane staining with anti-CD45 antibody in yellow. White arrows indicate intracellular bacteria, cyan arrows indicate extracellular bacteria. All scale bars, 10 μM. **(A.I,B.I,C.I,D.I)** Time-lapse microscopy image-series of macrophages that interacts with *Mtb* aggregates and fragment (A-C) or do not fragment (B-D) them. **(A.II,B.II,C.II,D.II)** Max intensity projection of confocal microscopy images of the same macrophages shown in panels I. **(A.III,B.III,C.III,D.III)** 3-D reconstruction of the cells imaged in panels II, images are cropped in x or y in the position indicated by the white dotted lines in panels II to show the inside of the cell.

**Supplementary Figure S5.**
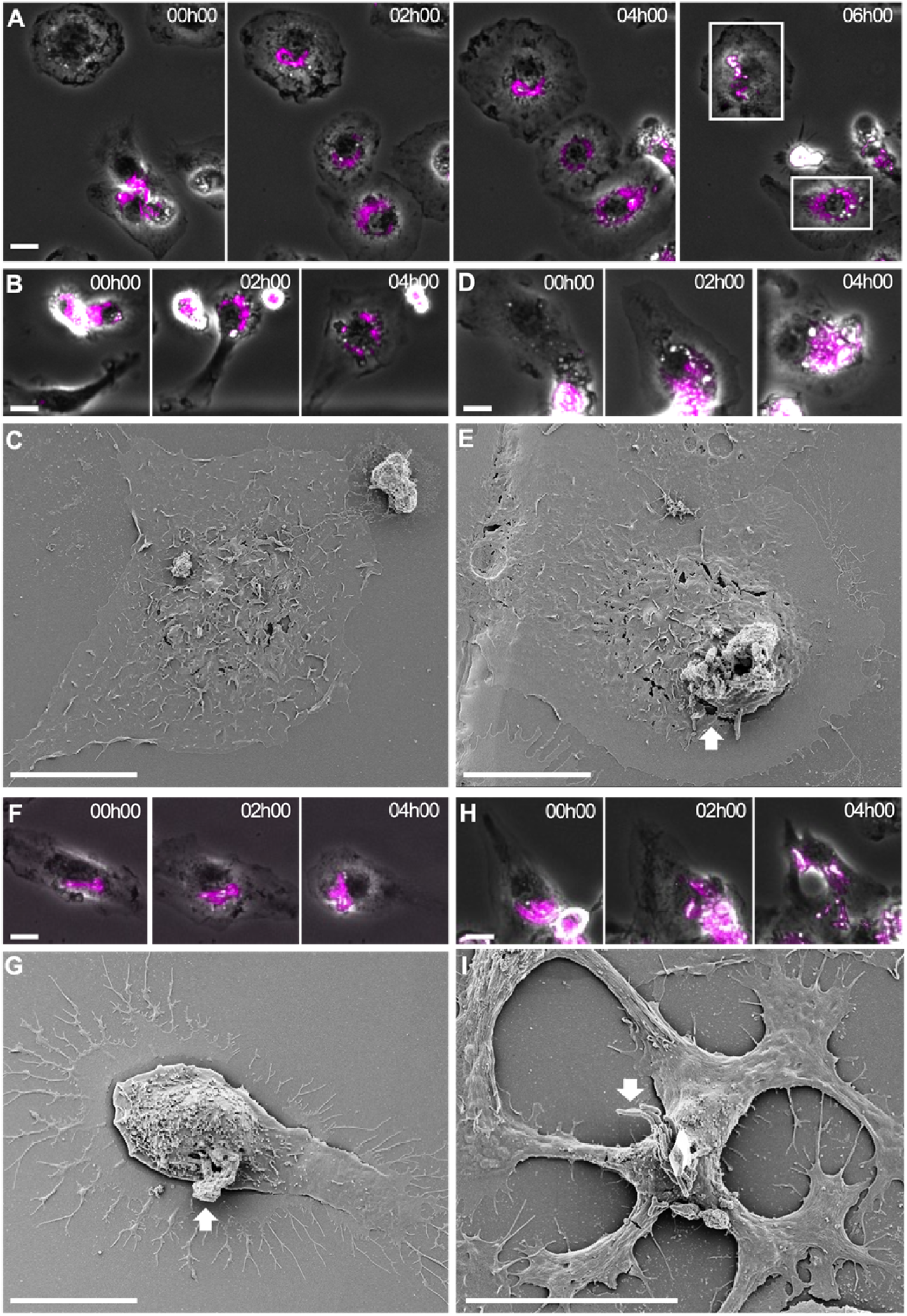
Correlative time-lapse microscopy and scanning electron microscopy (SEM) on BMDMs infected with *Mtb* aggregates. **(A-H)** BMDMs infected with aggregates of *Mtb* were imaged by time-lapse microscopy at 2-hour intervals for 12 hours, followed by SEM. **(A)** Time-lapse microscopy image-series of the macrophages shown in figure 1I,J (bottom square) and in figure 1K-M (top square). **(B)** Time-lapse microscopy image-series of a macrophage that shows the typical “bullseye” pattern of bacterial redistribution upon interaction with a bacterial aggregate. **(C)** Correlative SEM image of the macrophage shown in (B). **(D,F)** Time-lapse microscopy image-series of macrophages interacting with extracellular *Mtb* aggregates without fragmentation and redistribution of bacteria. **(E,G)** Correlative SEM images of (D) and (F) respectively. **(H)** Time-lapse microscopy image-series of a macrophage that interacts with an extracellular *Mtb* aggregates and fragments it. **(I)** Correlative SEM image of (H). This example shows that even some aggregates that get fragmented can be partially retained on the surface of the macrophage. Scale bars, 20 μm.

**Supplementary Figure S6.**
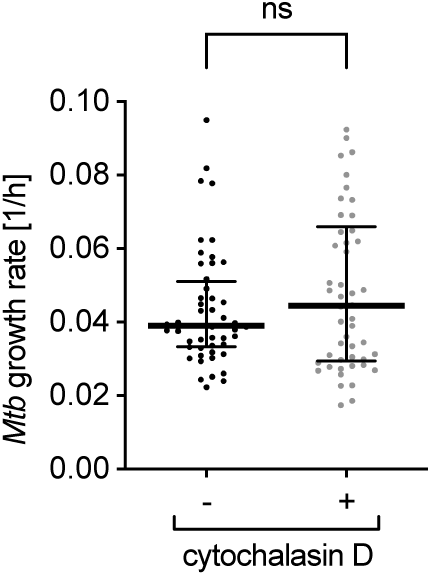
Cytochalasin D does not affect the growth of *Mtb* aggregates. BMDMs infected with aggregates of fluorescent *Mtb* and incubated without (-) or with (+) cytochalasin D were imaged by time-lapse microscopy at 1-hour intervals for 72 hours. The bacterial growth rate was calculated from microscopy time-series by measuring the fluorescent area over time of individual *Mtb* microcolonies. Each symbol represents an *Mtb* microcolony (n > 50 bacterial aggregates per condition). Black lines indicate median values and interquartile ranges. *P*-values were calculated using an unpaired Mann-Whitney test; ns, *P* values > 0.05.

**Supplementary Figure S7.**
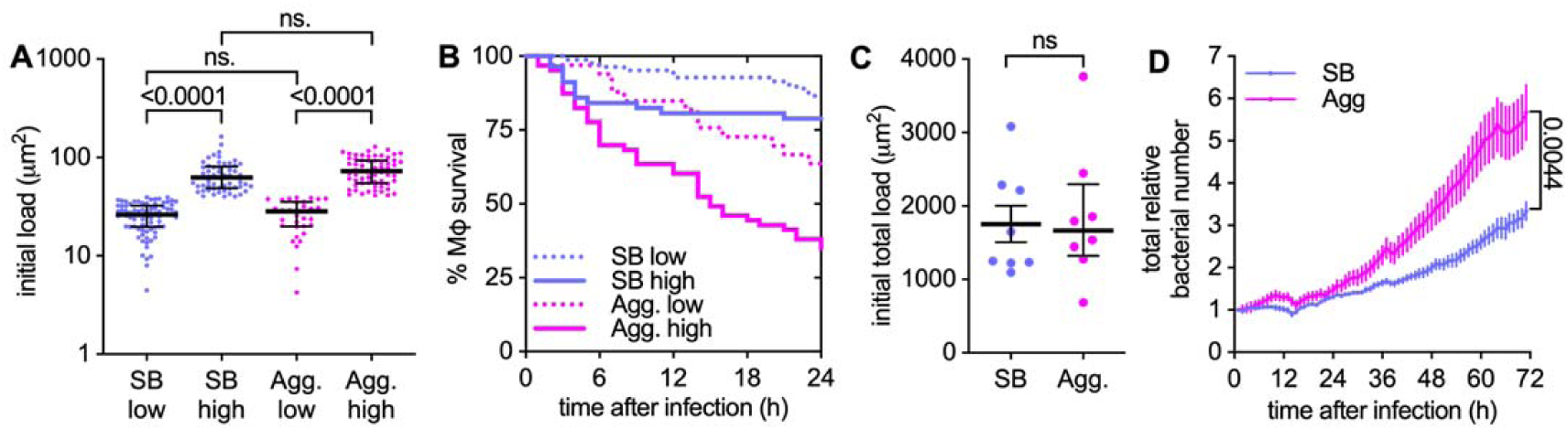
Bacterial aggregation enhances uptake-dependent killing of macrophages and bacteria propagation. **(A-D)** BMDMs were infected with aggregated (Agg) or non-aggregated (SB) *Mtb* Erd-tdTomato and imaged by time-lapse microscopy at 1-hour intervals for 72 hours. **(A)** Infected individual macrophages are binned into low and high initial load according to the amount of single (SB) or aggregated (Agg) bacteria they internalize. The bacterial load is calculated as fluorescent area per macrophage and gates are set at < 40 μm^2^ (low) or > 40.0 μm^2^ (high) per macrophage. The area of one bacterium is included between 0.5 and 2 μm^2^. Each symbol represents the bacterial load of one individual macrophage (n= 82, 57, 33, 63 respectively). Bars represent median and interquartile range. P-value were calculated using a Krustal-Wallis test; ns, *P* values > 0.05. **(B)** Percentage survival over time for individual macrophages with an initial bacterial load as indicated in panel A. **(C)** Total initial bacterial load per microscopy field of view (332.80 x 332.80 μm^2^, approx. 100 cells/field of view). Each symbol represents one field of view (n=8). Bars represent mean and standard errors of the mean. P-value calculated using an unpaired t test; *P* values > 0.05. **(D)** Total relative bacterial load over time per microscopy field of view. Symbols represent the average bacterial load (n=8), bars represent standard errors of the mean. P-value were calculated using an unpaired t test.

**Supplementary Figure S8.**
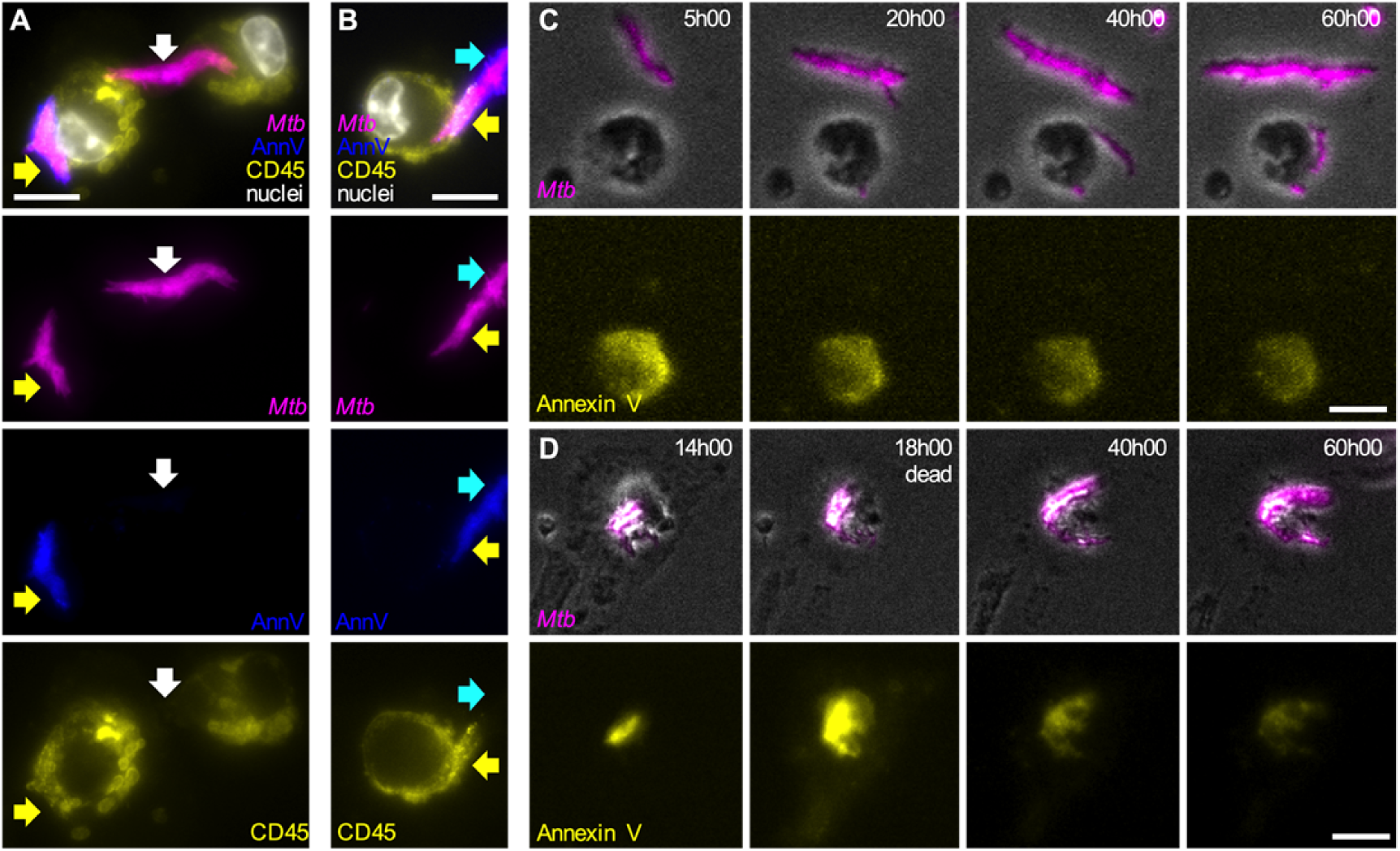
Formation of Annexin V-positive membrane domains requires physical contact between *Mtb* aggregates and live macrophages. **(A,B)** Representative fluorescence microscopy images of cytochalasin D-treated BMDMs infected with aggregates of *Mtb* Erd*-*tdTomato in the presence of Annexin V-FITC and fixed at 8 hours post infection. The plasma membrane of the cells was stained with and anti CD-45 antibody and nuclei were stained with Hoechst (white). Yellow arrows point at *Mtb* aggregates (magenta) overlapping with areas that stain positive for Annexin V (blue) and macrophages plasma membrane (yellow). White arrows indicate an *Mtb* aggregate that do not colocalize neither with the macrophages plasma membrane now with an Annexin V area. Cyan arrows indicate the distal area of an *Mtb* aggregate that stains positive for Annexin V but does not colocalize with the macrophages plasma membrane. Scale bars, 20 μm. **(C,D)** BMDMs treated with cytochalasin D were infected with aggregates of *Mtb*, incubated with Annexin V and imaged by time-lapse microscopy at 1-hour intervals for 60 hours. **(C)** Example of bacterial aggregates (magenta, top panels) that do not interact with macrophages and never become Annexin V-positive (yellow, bottom panel) during the course of the experiment. **(D)** Example of a bacterial aggregates (magenta, top panels) that induces formation of local an Annexin V-positive membrane domain in the interacting macrophage (yellow, bottom panel). After death of the macrophage (at 18h00) the bacterial aggregates gradually lose fluorescence (40h00-60h00). Scale bars, 20 μm.

**Supplementary Figure S9.**
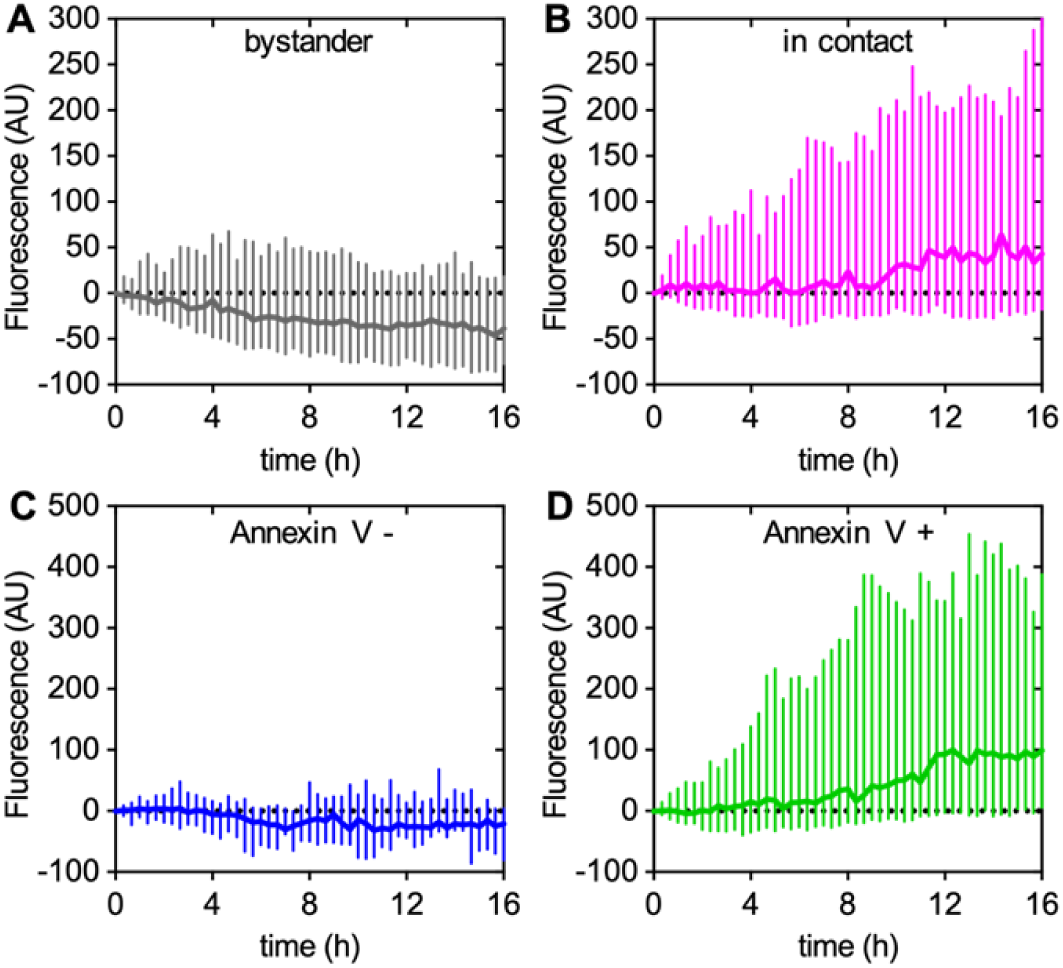
Extracellular *Mtb* aggregates induce calcium accumulation over time in cytochalasin D-treated macrophages. BMDMs stained with Oregon Green 488 Bapta-1 AM and treated with cytochalasin D were infected with aggregates of *Mtb* Erd-tdTomato and imaged by time-lapse microscopy at 20-minute intervals for 24 hours. Oregon Green 488 Bapta-1 AM fluorescence values at each time point were normalized to the time of first contact for infected cells or to time 0 for uninfected bystander cells. Lines represent median fluorescence values for all cells, error bars represent interquartile ranges. **(A,B)** Cells that survived until the end of the experiment and that are bystander (**A**, n=88) or in stable contact with *Mtb* aggregates (**B**, n=67). The distributions of the fluorescence values at 16 h in A and B are significantly different, *p* value = 0.0009, calculated using a Welch’s t test. **(C,D)** Cells in stable contact with *Mtb* aggregates that do not (**C**, n=24) or do (**D**, n=65) develop Annexin V-positive membrane domains. Both the cells that stay alive and that die during the course of the experiment are included in the plots. The distributions of the fluorescence values at 16 h in C and D are significantly different, *p* value < 0.0001, calculated using a Welch’s t test.

**Supplementary Figure S10.**
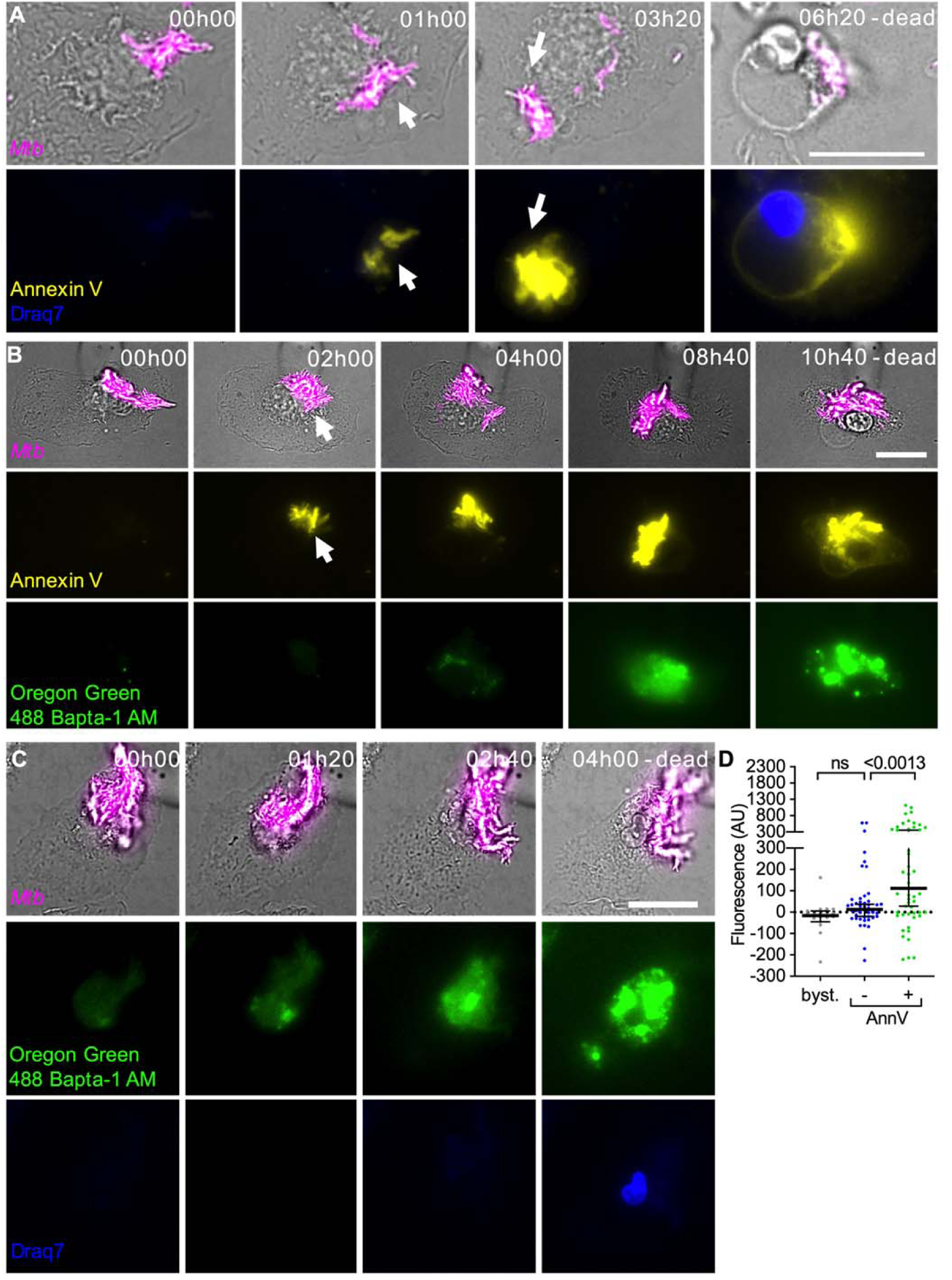
Extracellular *Mtb* aggregates induce formation of Annexin V-positive plasma membrane domains, intracellular calcium accumulation, and death in macrophages not treated with cytochalasin D. BMDMs were infected with aggregates of *Mtb* Erd-tdTomato and imaged by time-lapse microscopy at 20-minute intervals for 24 hours. Scale bars, 20 μm. **(A)** Example of a BMDM (brightfield) interacting with an extracellular *Mtb* aggregate (magenta). Annexin V-positive plasma membrane domains (yellow). Nucleus of dead cell stained with Draq7 (blue). White arrows indicate co-localization between bacteria and a local Annexin V-positive plasma membrane domain. **(B)** Example of a macrophage (brightfield) interacting with an extracellular *Mtb* aggregate (magenta). Annexin V-positive plasma membrane domains (yellow). Cytoplasmic Ca^2+^ stained with Oregon Green 488 Bapta-1 AM (green). Annexin V and Oregon Green 488 Bapta-1 AM staining increase over time until cell death at 10h40 after first contact. **(C)** Example of a macrophage (brightfield) interacting with an extracellular *Mtb* aggregate (magenta). Cytoplasmic Ca^2+^ stained with Oregon Green 488 Bapta-1 AM (green). Nucleus of dead cell stained with Draq7 (blue). Oregon Green 488 Bapta-1 AM staining increases over time until cell death at 4h00 after first contact. **(D)** Oregon Green 488 Bapta-1 AM fluorescence in uninfected bystander cells and for infected cells with (+) or without (–) Annexin V-positive plasma membrane domains at the site of contact with an *Mtb* aggregate. Oregon Green 488 Bapta-1 AM fluorescence values for infected cells correspond to the time of death after first contact or 16 hours post-contact for cells that survive. Values for uninfected bystander cells correspond to the time of death or 16 hours. Each symbol represents a single macrophage. Black bars represent median and interquartile range. *P*-values were calculated using a Krustal-Wallis test (n = 18, 46, 52 respectively). ns, *P* values > 0.05.

**Supplementary Figure S11.**
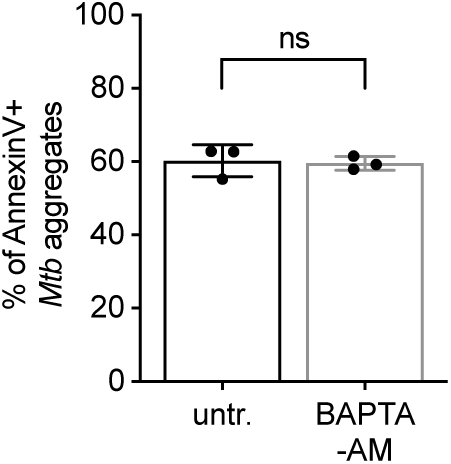
Calcium chelation does not affect formation of Annexin V-positive local membrane domains in macrophages in contact with *Mtb* aggregates. BMDMs treated with cytochalasin D were infected with aggregates of *Mtb*, incubated with Annexin V without or with BAPTA-AM and imaged by time-lapse microscopy at 1-hour intervals for 60 hours. The plot represents the percentage of macrophages that show Annexin V-positive membrane domains within the first 12 hours after entering in contact with *Mtb* aggregates. Each symbol represents a single experimental replicate. At least 100 individual macrophage-*Mtb* aggregate interactions were tracked for each replicate. Bars represent average and standard deviation. *P*-values were calculated using an unpaired t test; ns, *P* values > 0.05.

**Supplementary Figure S12.**
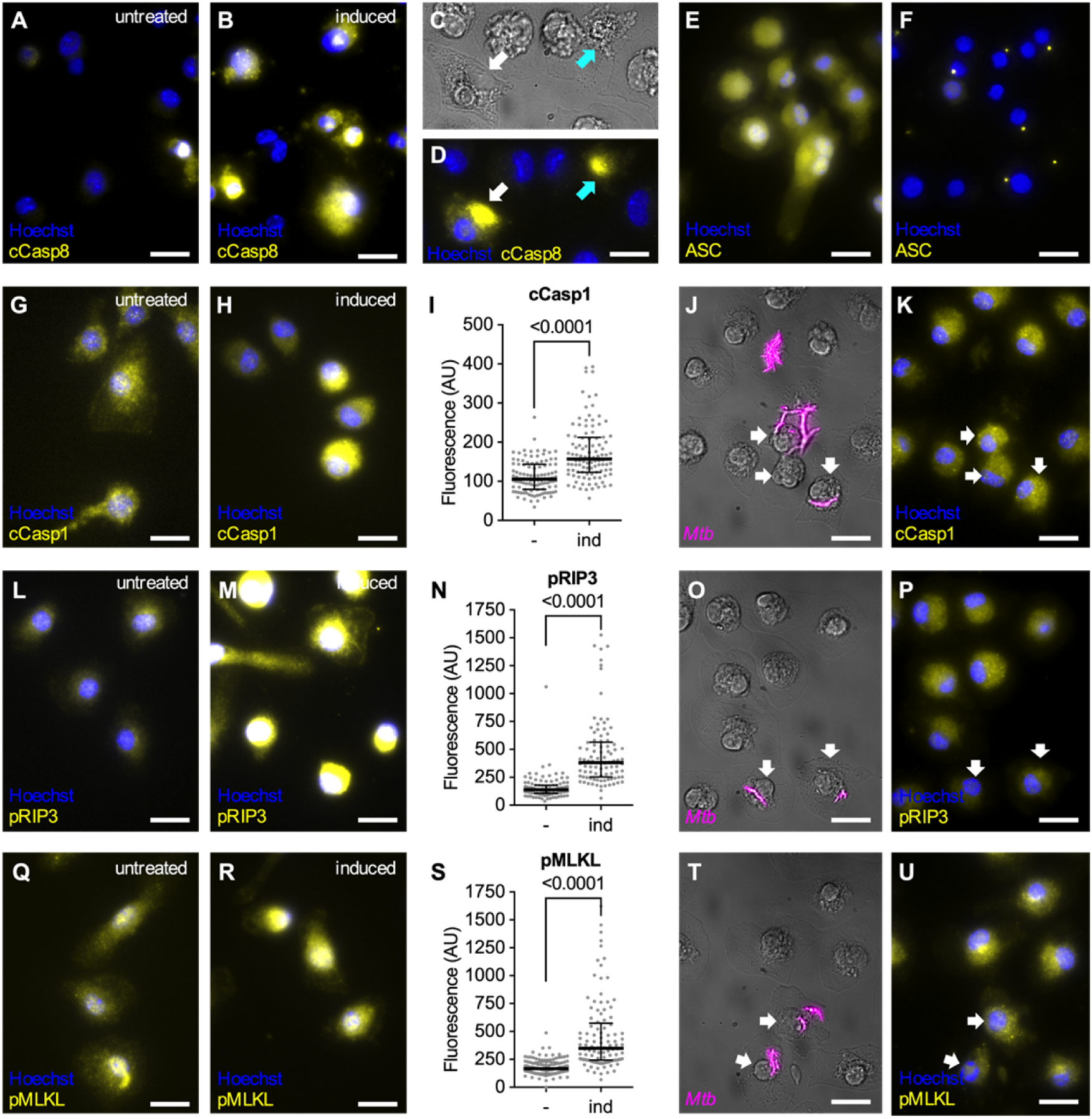
Apoptosis, pyroptosis and necroptosis markers can be induced chemically, but only pyroptosis markers are induced by contact with *Mtb.* **(A,B,E-I,L-N,Q-S)** Cytochalasin D-treated BMDMs were incubated without **(A,E,G,I,L,N,Q,S)** or with apoptosis inducers (20 ng/ml TNFα, cycloheximide 10 μg/ml) (31) **(B)**, pyroptosis inducers (50 ng/ml LPS, 0.5 mM ATP) (91) **(F,H,I)** or necroptosis inducers (5 μM zVAD, 20 ng/ml TNFα, 50 nM SM164) (31) **(M,N,R,S)** for 2 hours, fixed and processed for immunofluorescence with a anti-cleaved Caspase-8 antibody (cCasp8) **(A,B)**, a anti-ASC antibody **(E,F)**, a anti-cleaved Caspase-1 antibody (cCasp1) **(G-I)**, a anti-phosphorylated RIP3 antibody (pRIP3) **(L-N)** or a anti-phosphorylated MLKL antibody (pMLKL) **(Q-S)**. **(A,B,E,F,G,H,L,M,Q,R)** Representative microscopy images of BMDMs treated without (left panels-untreated) or with (right panels-induced) death pathways inducers. Antibody staining in yellow, nuclei in blue (stained with Hoechst). Scale bars, 10 μm. **(I,N,S)** Median fluorescence values for BMDMs treated without (-) or with (ind) death pathways inducers. Each symbol represents a single macrophage. Black bars represent median and interquartile range. P-value were calculated using an unpaired Mann-Whitney test. (**I:** n=108, 109 respectively; **N:** n=103, 94 respectively; **S:** n=110, 109 respectively). **(C,D,J,K,O,P,T,U)** Representative microscopy images of cytochalasin D-treated BMDMs infected with aggregates of *Mtb* Erd-tdTomato, fixed at 24 hours post infection and processed for immunofluorescence with a anti-cleaved Caspase-8 antibody (cCasp8) **(C,D)**, a anti-cleaved Caspase-1 antibody (cCasp1) **(J,K)**, a anti-phosphorylated RIP3 antibody (pRIP3) **(O,P)** or a anti-phosphorylated MLKL antibody (pMLKL) **(T,U)**. White arrows indicate the cells that are considered as “in contact” for quantitative fluorescence analysis, the other cells are considered as “bystander”. **(C,J,O,T)** Representative microscopy images showing the cells (brightfield) overlapped with the fluorescence channel corresponding to the bacteria (magenta). **(D,K,P,U)** Fluorescence channels showing the indicated antibody staining (yellow) and the nuclei (blue, stained with Hoechst). Scale bars, 10 μm. In **C** and **D** we show that both dying cells with intact nuclei (white arrow) and dead cells with disrupted nuclei (cyan arrow) show signs of Caspase-8 activation.

**Supplementary Figure S13.**
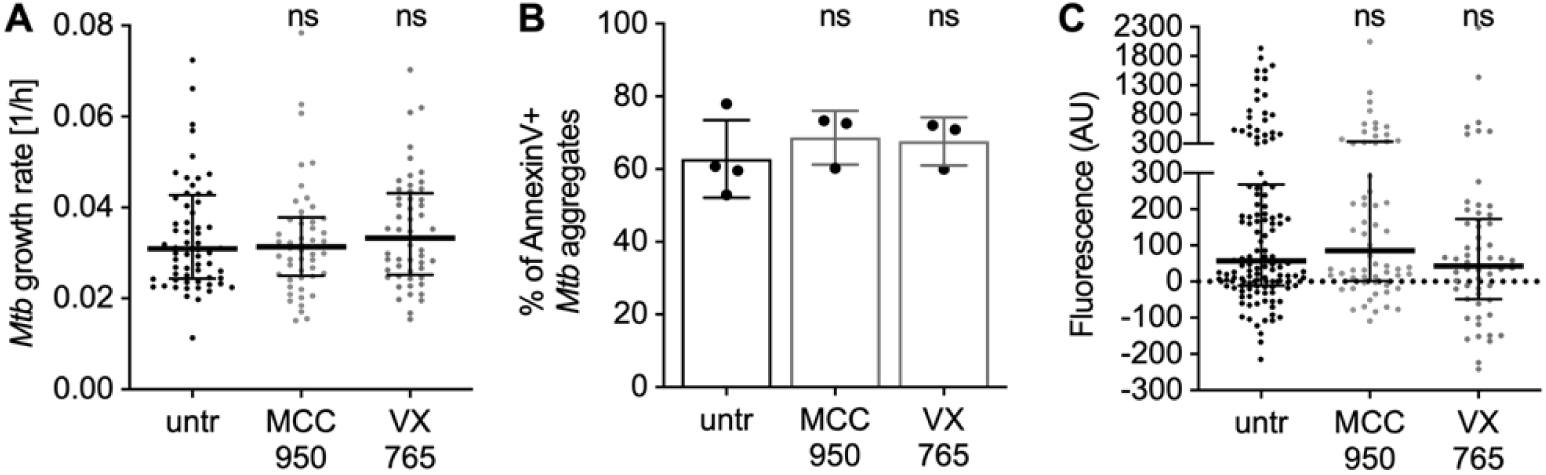
Treatment with pyroptosis inhibitors does not affect *Mtb* growth, formation of Annexin V-positive local membrane domains and intracellular calcium accumulation in macrophages in contact with *Mtb* aggregates. **(A)** BMDMs treated with cytochalasin D were infected with aggregates of *Mtb,* incubated with the indicated pyroptosis inhibitors and imaged by time-lapse microscopy at 1-hour intervals for 60 hours. The bacterial growth rate was calculated from microscopy time-series by measuring the fluorescent area over time of individual Mtb microcolonies. Each symbol represents an Mtb microcolony (n ≥ 50 bacterial aggregates per condition). Black lines indicate median values and interquartile ranges. P-values were calculated using a Krustal-Wallis test; ns, *P* values > 0.05. **(B)** BMDMs treated with cytochalasin D were infected with aggregates of *Mtb*, incubated with Annexin V and the indicated pyroptosis inhibitors and imaged by time-lapse microscopy at 1-hour intervals for 60 hours. The plot represents the percentage of macrophages that show Annexin V-positive membrane domains within the first 12 hours after entering in contact with *Mtb* aggregates. Each symbol represents a single experimental replicate. At least 70 individual macrophage-*Mtb* aggregate interactions were tracked for each replicate. Bars represent average and standard deviation. *P* values were calculated using a one-way ANOVA test comparing treated samples with the untreated control; ns, *P* values > 0.05. **(C)** Cytochalasin D-treated BMDMs were stained with the membrane-permeable dye Oregon Green 488 Bapta-1 AM to visualize cytoplasmic Ca^2+^, infected with aggregates of *Mtb* and incubated with the indicated death inhibitors. Samples were imaged by time-lapse microscopy at 20-minute intervals for 24 hours. Oregon Green 488 Bapta-1 AM fluorescence values at each time point were normalized to the time of first contact with an *Mtb* aggregate. Plotted values represent relative Oregon Green 488 Bapta-1 AM fluorescence values at the time of death (or to 16 hours post-contact for cells that survive) in untreated (untr.; n=124) infected macrophages and in infected macrophages treated with MCC950 (n=64) or VX765 (n=62). Each symbol represents a single macrophage. Black bars represent median and interquartile range. *P*-values were calculated using a Krustal-Wallis test comparing treated samples with the untreated control; ns, *P* values > 0.05.

**Supplementary Figure S14.**
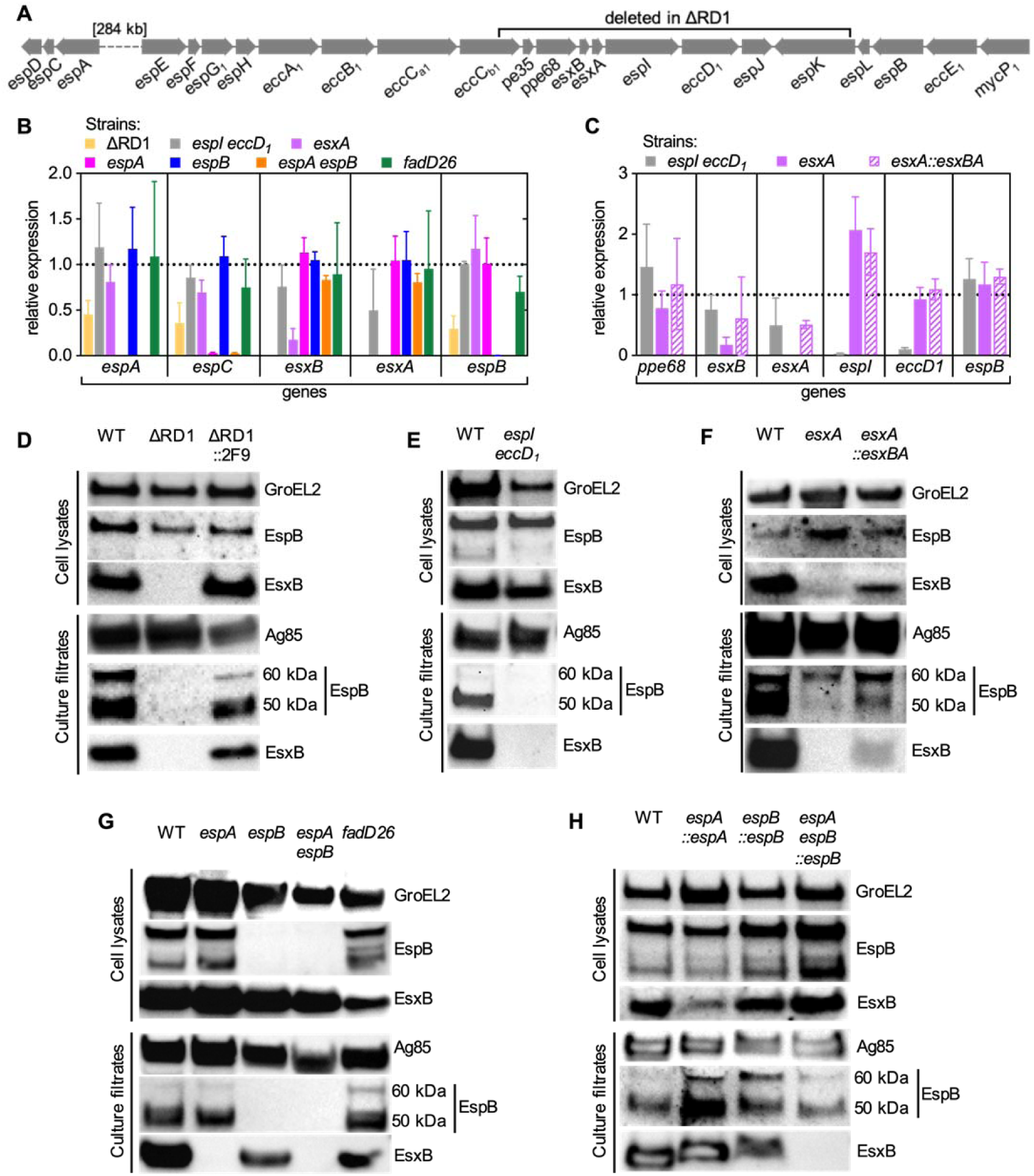
Expression and secretion patterns of ESX-1 proteins in the different mutant strains used in this study. **(A)** Representation of the *espACD* and *esx-1* loci in the *Mtb* genome. **(B-C)** Expression levels of selected genes of the *espACD* and *esx-1* loci in different mutants. Relative expression (fold changes) was normalized to the WT strain. **(D-H)** Representative immunoblots of cell lysates and culture filtrates from different *Mtb* strains probed with the indicated antibodies. In culture filtrates EspB shows a full-length 60-kDa isoform and a truncated 50-kDa isoform as previously reported (52, 57). GroEL2 and Ag85 were respectively used as loading controls for cell lysates and culture filtrates.

**Supplementary Figure S15.**
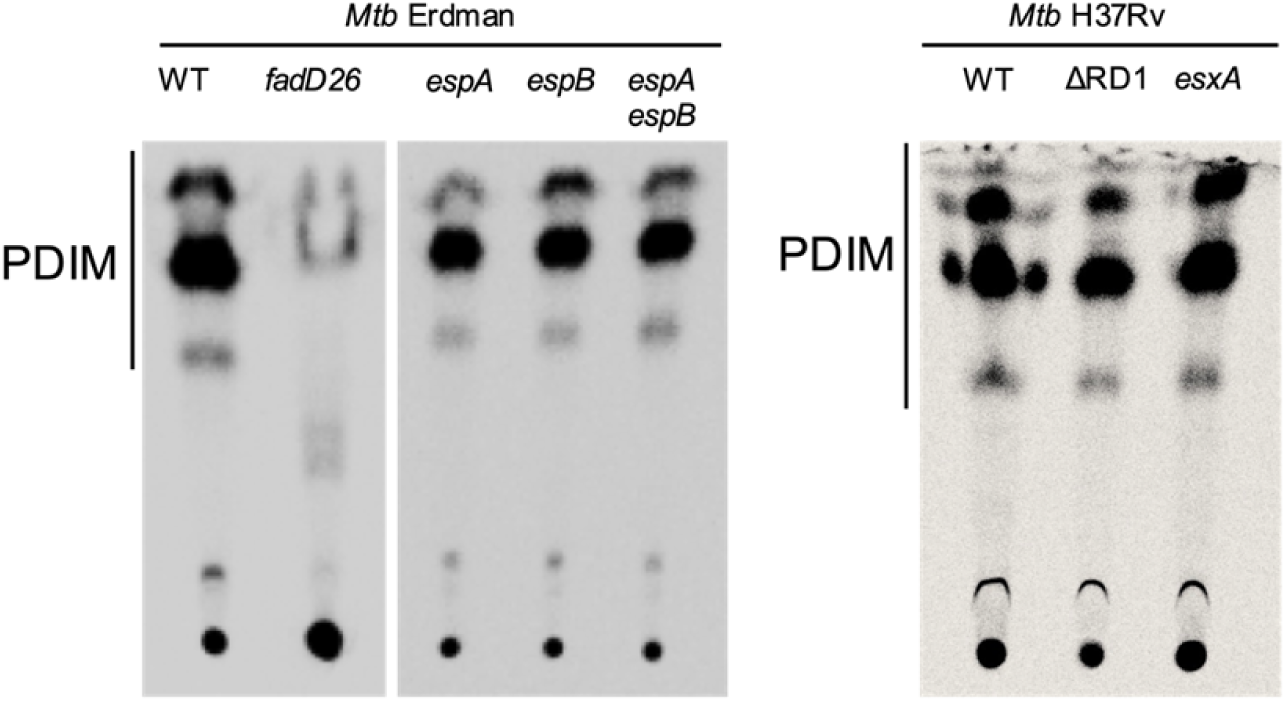
TLC analysis of PDIM production by wild-type and mutant *Mtb* strains used in this study. C^14^-labelled surface-exposed lipids were collected from *Mtb* pellets by petroleum ether extraction and spotted on a TLC silica gel plate. The plate was resolved in a mobile phase of 9:1 petroleum ether-diethyl ether.

**Figure S16.**
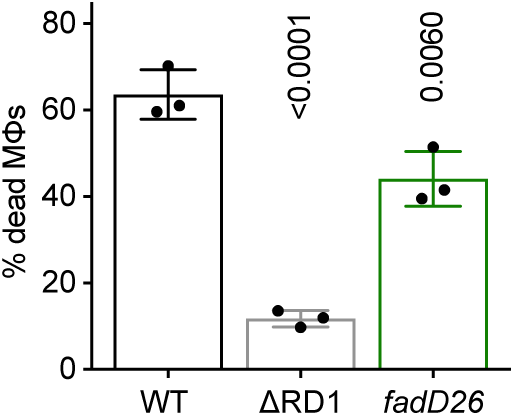
Lack of PDIM has a minor effect on uptake-dependent killing of macrophages by *Mtb* aggregates. BMDMs were infected with aggregates of different *Mtb* strains and imaged by time-lapse microscopy at 1-hour intervals for 60 hours. The plots represent the percentage of macrophages that die within the first 12 hours after interaction with an *Mtb* aggregate. Each symbol represents the percentage of dead macrophages for a single experimental replicate (n ≥ 3 replicates with ≥ 70 cells per replicate). Bars represent means and standard deviations. *P*-values were calculated using a one-way ANOVA test.

**Supplementary Figure S17.**
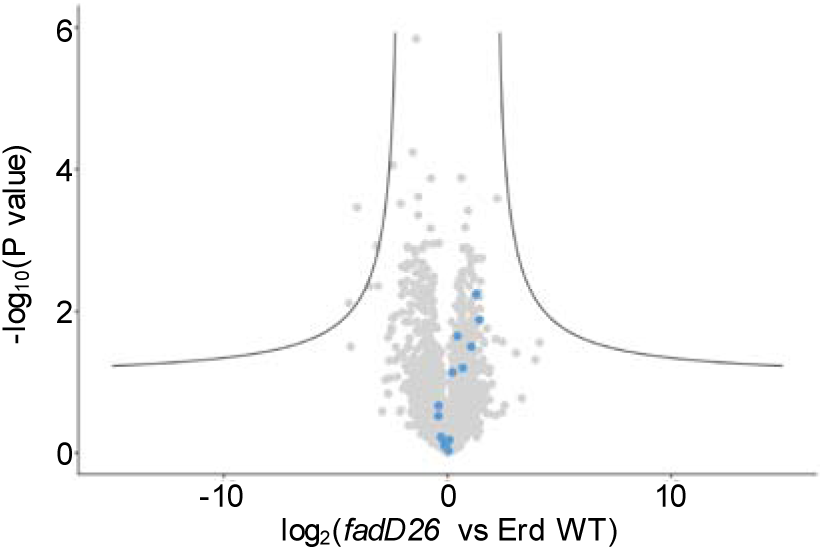
Mass spectrometry analysis of the bacterial secretome of the *fadD26* mutant. Volcano plot showing changes in secreted protein abundances in the *fadD26* mutant compared to the Erdman WT parental strain. Black lines correspond to a false discovery rate (FDR) of 0.05, S0=1. Light blue symbols represent proteins that are part of the ESX-1 operon.

**Supplementary Figure S18.**
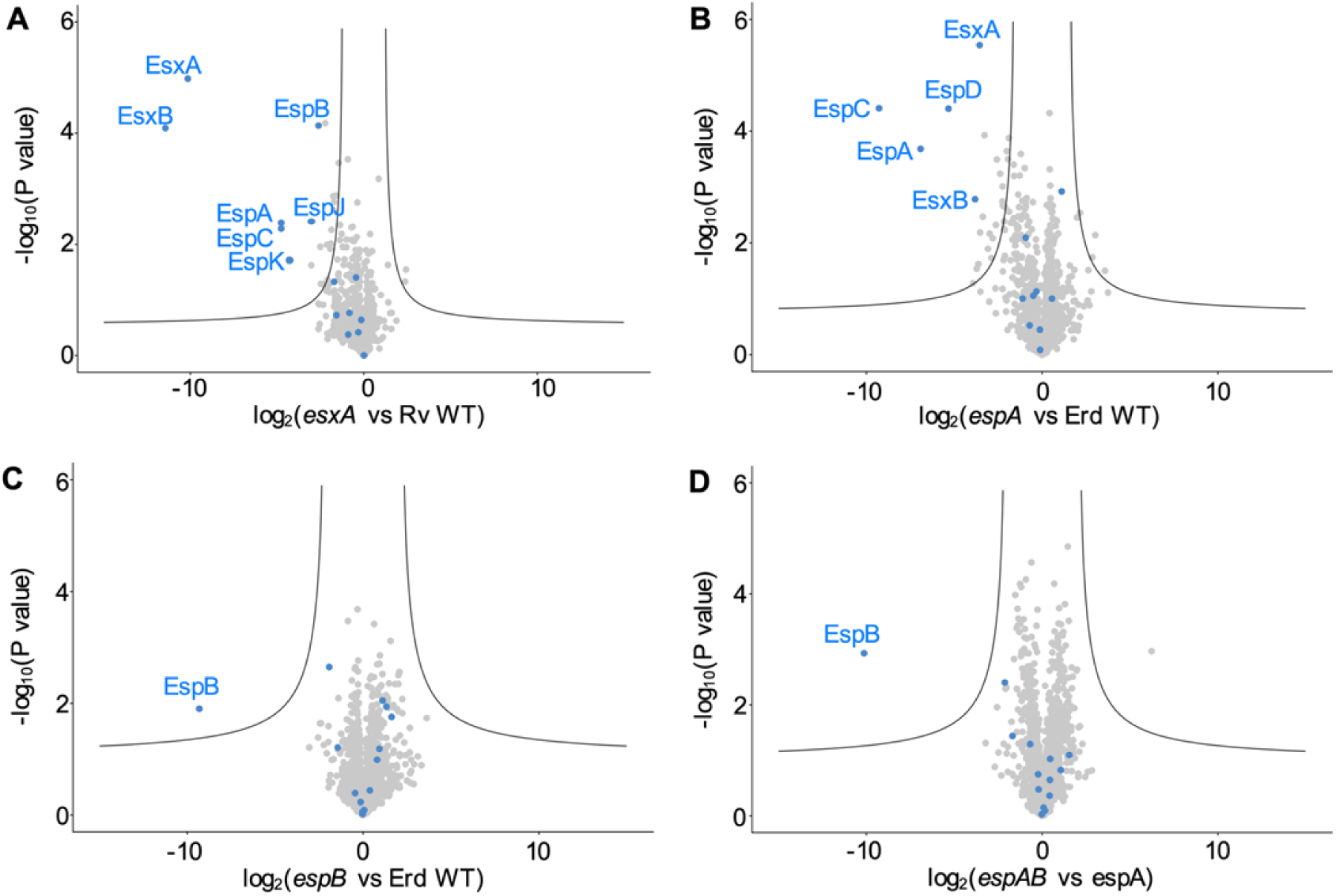
Mass spectrometry analysis of the bacterial secretome. Volcano plots showing changes in secreted protein abundances between couples of *Mtb* strains. Black lines correspond to a false discovery rate (FDR) of 0.05, S0=1. Light blue symbols represent proteins that are part of the ESX-1 operon. The names of the ESX-1 operon proteins showing a significant change are indicated. **(A)** *esxA* mutant strain compared to the Rv WT parental strain; **(B)** *espA* mutant strain compared to the Erdman WT parental strain; **(C)** *espB* mutant strain compared to the Erdman WT parental strain; **(D)** *espA espB* mutant strain compared to the *espA* mutant parental strain.

**Supplementary Figure S19.**
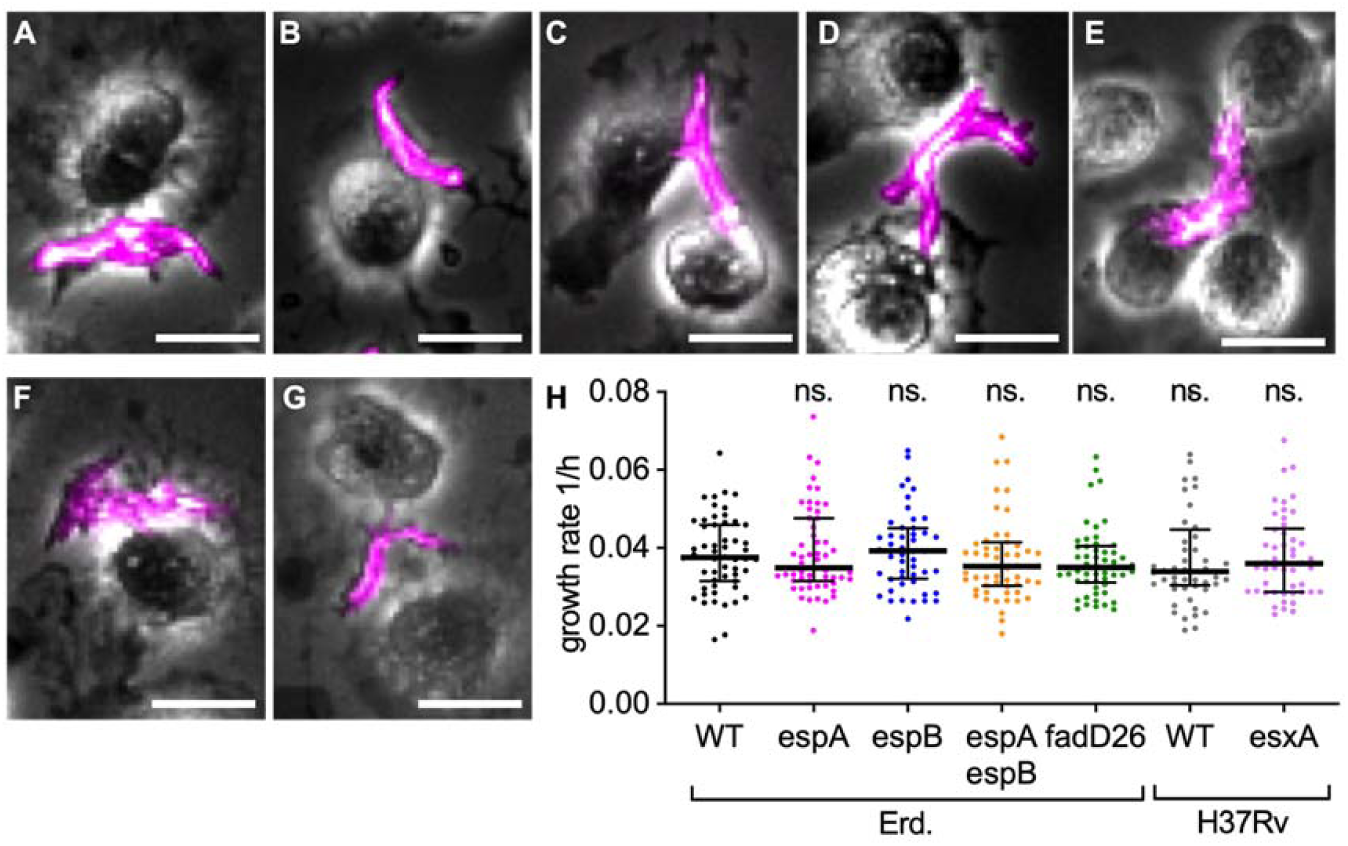
Bacterial aggregates from different *Mtb* strains have comparable morphology and show similar growth dynamics. **(A-G)** Representative examples of aggregates of different fluorescent *Mtb* strains in contact with cytochalasin D-treated BMDMs at 5 hours post infection. Scale bars, 20 μm. **(A)** *Mtb* Erdman WT expressing tdTomato; **(B)** *Mtb espA* expressing tdTomato; **(C)** *Mtb espB* expressing tdTomato; **(D)** *Mtb espA espB* expressing tdTomato; **(E)** *Mtb fadD26* expressing tdTomato; **(F)** *Mtb* H37Rv WT expressing GFP; **(G)** *Mtb esxA* expressing GFP. **(H)** The bacterial growth rate was calculated from microscopy time-series by measuring the fluorescent area over time of individual *Mtb* microcolonies. Each symbol represents an *Mtb* microcolony (n ≥ 50 bacterial aggregates per condition). Black lines indicate median values and interquartile ranges. *P*-values were calculated using a Krustal-Wallis test comparing each strain with Erdman (Erd.) WT (black p-values); ns, *P* values > 0.05.

**Supplementary Figure S20.**
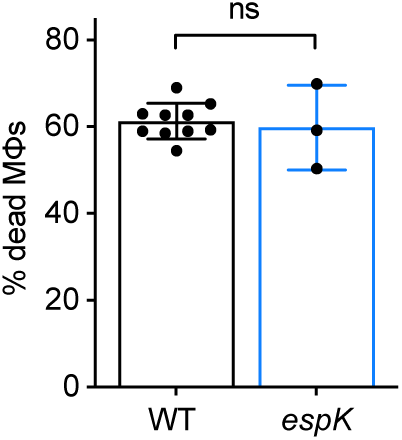
EspK is not required for uptake-independent killing of macrophages by *Mtb* aggregates. BMDMs treated with cytochalasin D were infected with aggregates of wild-type *Mtb*, or with aggregates of the *espK* mutant and imaged by time-lapse microscopy at 1-hour intervals for 60 hours. Plots represent the percentage of BMDMs that die within the first 12 hours after stable contact with an aggregate of *Mtb.* Each symbol represents the percentage of dead macrophages for a single experimental replicate (n ≥ 3 replicates with ≥ 70 cells per replicate). Bars represent means and standard deviations. *P*-values were calculated using an unpaired t test comparing the mutant strain to the wild-type control; ns, *P* values > 0.05.

**Supplementary Figure S21.**
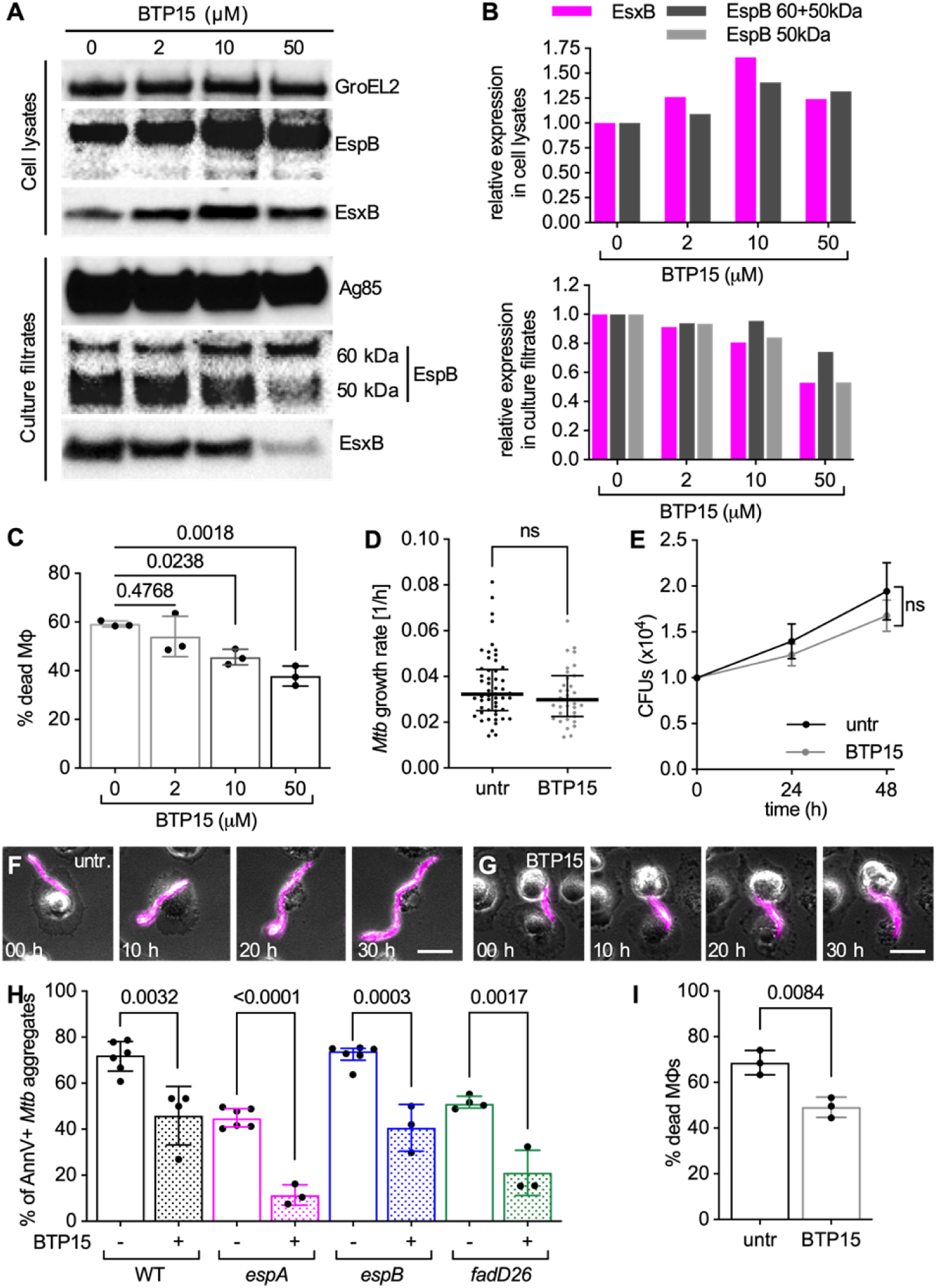
BTP15 treatment reduces ESX-1 secretion and local plasma membrane in macrophages in contact with *Mtb,* without affecting ESX-1 expression, aggregates growth dynamics and morphology in *Mtb*. **(A)** Western blot showing EspB and EsxB expression (cell lysates) and secretion (culture filtrates) pattern of *Mtb* Erdman WT cultures treated with different concentrations of BTP15 (0, 2, 10, 50 μM). GroEL2 and Ag85 were respectively used as loading controls for cell lysates and culture filtrates. **(B)** EspB (whole or 50 kDa isoform) and EsxB quantification from the Western blot images. Values were normalized to the loading control first and then to the untreated samples. **(C-H)** BMDMs treated with cytochalasin D were infected with aggregates of *Mtb* and incubated with or without BTP15. Infected cells were imaged by time-lapse microscopy at 1-hour intervals for 60 hours **(C,D,F-H)**, or incubated at 37°C with 5% CO_2_ for quantification of colony-forming units (CFU) at 0, 24, and 48 hours post-infection **(E)**. **(C)** Percentage of macrophages that die within the first 12 hours after interaction with an *Mtb* aggregate. Bacteria are incubated with different concentration of BTP15 (0, 2, 10, 50 μM) for 48 h before infection. BTP15 is not added to the medium of the cells during the course of the experiment. Each symbol represents a single experimental replicate. At least 70 individual macrophage-*Mtb* aggregate interactions were tracked for each replicate. Bars represent average and standard deviation. *P*-values were calculated using a one-way ANOVA test. **(D)** The growth rate of individual untreated and BTP15-treated (50 μM) *Mtb* aggregates was calculated by measuring the fluorescent area over time. Each symbol represents a microcolony. Black lines indicate median values and interquartile ranges (n ≥ 50 bacterial aggregates per condition). *P*-value calculated using an unpaired Mann-Whitney test; ns, *P* value > 0.05. **(E)** Total CFU per well at different time points from untreated and BTP15-treated (50 μM) bacterial cultures. Symbols and bars represent means and standards deviations (n = 4 biological replicates). *P*-value calculated using an unpaired t test; ns, *P* value > 0.05. **(F-G)** Representative examples of untreated (**F**) or BTP15-treated (50 μM) (**G**) aggregates of fluorescent *Mtb* in contact with cytochalasin D-treated BMDMs at different time-points post infection. Scale bars, 20 μm. **(H)** BMDMs treated with cytochalasin D were infected with aggregates of different *Mtb* strains, incubated with Annexin V and imaged by time-lapse microscopy at 1-hour intervals for 60 hours. Percentage of macrophages that show Annexin V - positive membrane domains within the first 12 hours after entering in contact with *Mtb* aggregates without (-) or with (+) BTP15 treatment (50 μM). Each symbol represents a single experimental replicate. At least 90 individual macrophage-*Mtb* aggregate interactions were tracked for each replicate. Bars represent average and standard deviation. *P*-values were calculated using an unpaired t test comparing the treated samples with their untreated reference. **(I)** BMDMs were infected with aggregates of *Mtb* strains, incubated without (untr) or with (BTP15) 50 μM BTP15 and imaged by time-lapse microscopy at 1-hour intervals for 60 hours. Percentage of macrophages that die within the first 12 hours after interaction with an *Mtb* aggregate. Each symbol represents a single experimental replicate. At least 100 individual macrophage-*Mtb* aggregate interactions were tracked for each replicate. Bars represent average and standard deviation. *P*-values were calculated using an unpaired t test.

**Table S1.**
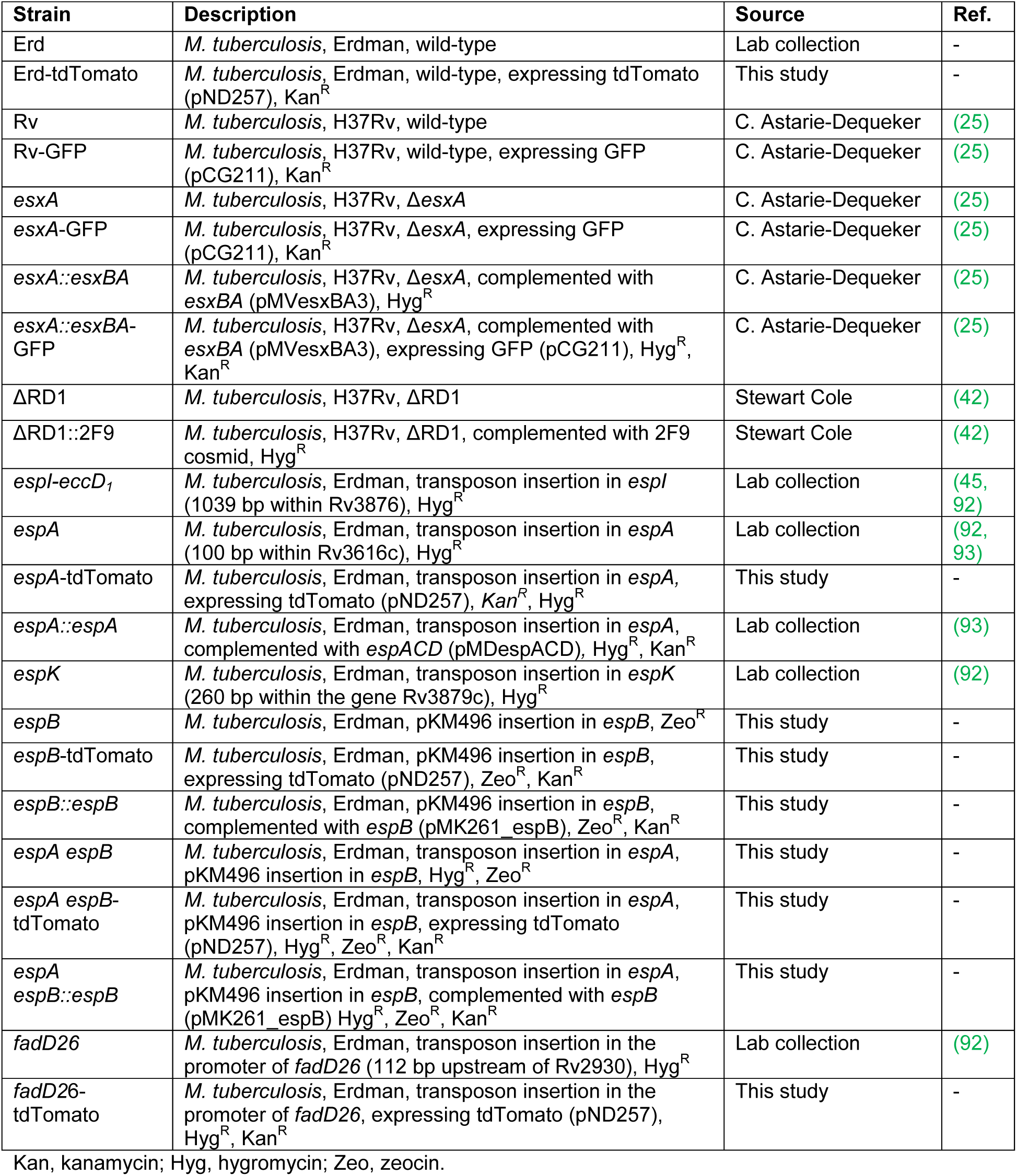
List of *Mtb* strains used in this study.

**Table S2.**
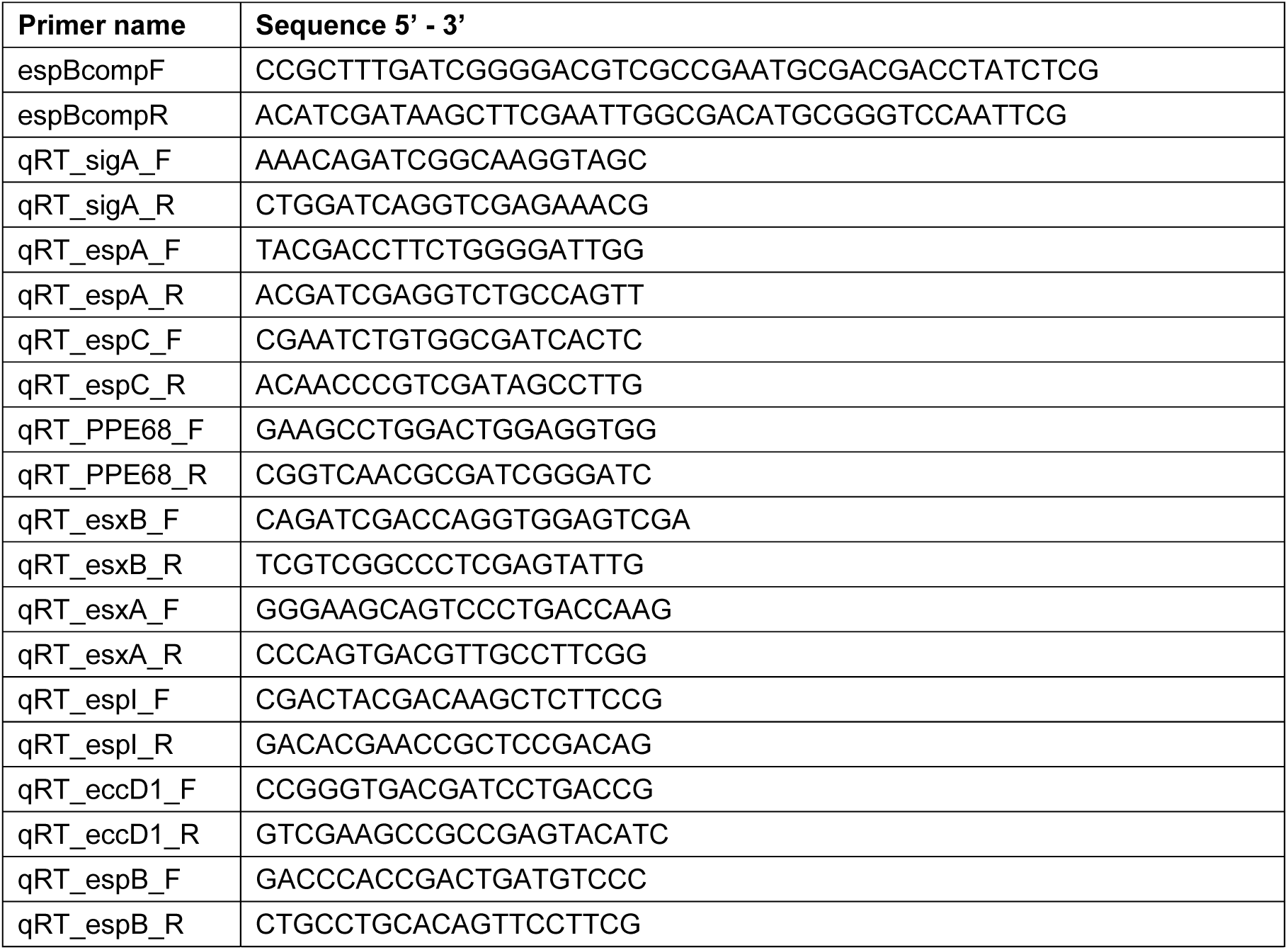
List of primers used in this study.

